# TarDis: Achieving Robust and Structured Disentanglement of Multiple Covariates

**DOI:** 10.1101/2024.06.20.599903

**Authors:** Kemal Inecik, Aleyna Kara, Antony Rose, Muzlifah Haniffa, Fabian J. Theis

## Abstract

Addressing challenges in domain invariance within single-cell genomics necessitates innovative strategies to manage the heterogeneity of multi-source datasets while maintaining the integrity of biological signals. We introduce *TarDis*, a novel deep generative model designed to disentangle intricate covariate structures across diverse biological datasets, distinguishing technical artifacts from true biological variations. By employing tailored covariate-specific loss components and a self-supervised approach, *TarDis* effectively generates multiple latent space representations that capture each continuous and categorical target covariate separately, along with unexplained variation. Our extensive evaluations demonstrate that *TarDis* outperforms existing methods in data integration, covariate disentanglement, and robust out-of-distribution predictions. The model’s capacity to produce interpretable and structured latent spaces, including its pioneering work in ordered latent representations for continuous covariates, markedly enhances its utility in hypothesis-driven research. Consequently, *TarDis* offers a promising analytical platform for advancing scientific discovery, providing insights into cellular dynamics, and enabling targeted therapeutic interventions.

**Progress and potential:** Modern single-cell genomics provides an unprecedented view into cellular heterogeneity, yet the very richness that propels new discoveries also complicates downstream analysis. Gene-expression patterns emerge from overlapping biological processes (e.g., differentiation programs, disease progression) and extrinsic factors (e.g., laboratory protocols, technical artifacts). *Disentanglement*, in this context, aims to parse these intertwined influences into interpretable latent representations, a crucial step for elucidating how complex covariates shape cellular states. While methods that correct for batch effects have become standard, these strategies often fall short in achieving the deeper objective of capturing subtle, high-dimensional biological dynamics. In single-cell experiments, cells navigate intricate developmental trajectories, respond nonlinearly to environmental or pharmaceutical perturbations, and exhibit myriad context-specific behaviors. Without disentanglement, these diverse signals frequently remain intermingled, limiting biological interpretability and hindering hypothesis-driven research.

Disentangling biological covariates is particularly vital for addressing nuanced questions in single-cell research. For example, in a disease model involving multiple genetic variants and variable drug dosing, researchers may wish to examine the effect of each variant independently or investigate how dosage influences a specific mutant background. Similarly, in developmental biology, uncovering how cells evolve across a continuum of pseudotime (e.g., from pluripotent to fully differentiated states) is critical for identifying the genes that orchestrate fate decisions while isolating the influence of developmental time from tissue-specific contexts, along with other confounding factors such as culture conditions, sample preparation, or donor genetic characteristics. Alternatively, disentangling lineage commitment signals from spatial patterning cues enables the identification of master regulators driving fate decisions. Moreover, by explicitly isolating and representing each covariate as an independent latent dimension, one can systematically navigate and interrogate a rich *multidimensional covariate space*. This approach extends beyond merely observing biological states, it enables exploration of novel or unmeasured cellular conditions through latent-space manipulations. For instance, disentangled latent spaces could allow researchers to computationally predict cellular responses at drug dosages or developmental stages that were never experimentally observed, significantly broadening the scope and predictive power of experimental datasets. Such analyses yield testable hypotheses for unexplored biological phenomena and enable informed planning of subsequent experimental validations.

The challenge of covariate disentanglement stems fundamentally from the complexity of modeling joint distributions of gene expression conditioned simultaneously on multiple covariates, both categorical (e.g., tissue type, disease condition) and continuous (e.g., pseudotime, dosage). This is inherently an underdetermined problem because single-cell measurements represent only sparse snapshots within a vast combinatorial space of covariate conditions. Conventional modeling approaches often conflate correlated covariates, collapsing biological variability into ambiguous latent factors, and typically fail to explicitly create separate latent representations for disentangled covariates. Moreover, continuous covariates introduce an additional layer of complexity; yet discretizing them artificially imposes arbitrary boundaries, obscuring subtle transitions and hindering accurate capture of biological gradients. Therefore, preserving the continuous nature of such covariates in disentangled representations is critical, as it maintains their intrinsic ordering and enables researchers to discern nuanced biological shifts—such as identifying thresholds in dose-response relationships or characterizing gradual developmental transitions—in a naturally interpretable manner.

The key idea in this paper is to devise a tailored deep generative model for systematically separating both categorical and continuous covariates into independent latent dimensions, while still ensuring coherent integration of the underlying gene-expression data. By explicitly *targeting* these covariates and preserving continuous variables as smooth, ordered latent axes, our approach clarifies complex interactions and uncovers nuanced patterns that remain concealed under standard analyses. The resulting disentangled representations can then support robust out-of-distribution generalizations, refined differential analyses, and more principled hypotheses about how diverse factors interact to drive cellular variation.

## 1 Introduction

Domain invariance tackles the challenge of learning from datasets that, while representing the same physical phenomena, originate from disparate sources such as different users, acquisition devices, or locations [5]. As the data source often lacks direct relevance to the task, the objective is to develop a model that maintains performance robustness by being invariant to these domain variations. This invariance not only enhances model reliability across shifts, whether subpopulational [47] or distributional [27], but also is an end in itself where the source is obscured to comply with data protection requirements [30]. Such shifts, frequently observed in practical machine learning scenarios, necessitate models to be resilient to variations in multi-domain datasets by learning to minimize the disparity in data distributions within the representation space; ideally achieving a low metric distance between them. This concept is closely aligned with distributionally robust optimization strategies, promoting the development of universally applicable machine learning models that withstand out-of-distribution variations [28, 58, 85, 99].

The identification of spurious correlations within these multi-domain datasets can provide critical insights for certain downstream applications, enriching the interpretive scope beyond mere domain invariance. Moreover, models leveraging data representations or predictors derived from true correlations, including domain-specific attributes or nuisance factors, more effectively discern causal relationships, thereby enhancing their generalization capabilities [3, 4]. This recognition has spurred interest in disentangled representation learning, aiming to segregate and independently model spurious and invariant characteristics within the data [4, 6, 48]. Developing invariant representation learning models is a complex multi-objective optimization problem, frequently necessitating linear constraints on the data representations and classifiers [2, 3, 48], or the incorporation of conditional priors within the VAE framework [4, 59].

Existing invariant representation learning methods often fail for continuous domain problems, an area that is significantly underexplored yet critically important [7, 100, 102]. Examples include patient monitoring systems where physiological spurious data varies daily and across activities [18], finance, where models predicting stock prices or market trends must generalize across varying economic conditions and times [37], and climate modeling, where models use invariant learning to forecast weather or long-term climate changes across diverse locations and time periods [10]. Existing methods are generally designed for discrete categorical domains and struggle with the continuous nature of many real-world tasks. This leads to challenges such as sparse data in each domain, making it hard to accurately estimate invariant correlations, and segmentation of continuous data into discrete blocks which can misrepresent true data distributions. Addressing these issues is crucial for advancing model robustness and ensuring applicability in dynamic environments [100].

In the context of domain invariance, multi-domain and multi-condition single-cell genomics datasets present a critical testbed where the integration of data representations confronts complex challenges in biological and pharmaceutical research [33]. Single-cell genomics offers a granular view of individual cells’ genetic diversity, highlighting the variability among cells and essential for understanding cellular and molecular processes [9, 40, 69]. However, the data often come from a range of labs and varied experimental setups, incorporating batch effects and technical artifacts that can mask true biological signals (Figure 1a) [24, 55]. These challenges are compounded when data includes cells affected by chemical or genetic perturbations, sourced from diseased states, or differing in their origin, such as specific organs, organisms, developmental stages, ethnicity, age, sex and other factors that further contribute to variability [31, 35, 65, 80, 84]. Effective data integration is vital for separating technical artifacts from relevant biological signals, facilitating a robust comparison of biological landscapes across various domains and enhancing our understanding of the underlying cellular dynamics, with significant implications for advancing disease research and therapeutic development [72, 73].

**Figure 1.**
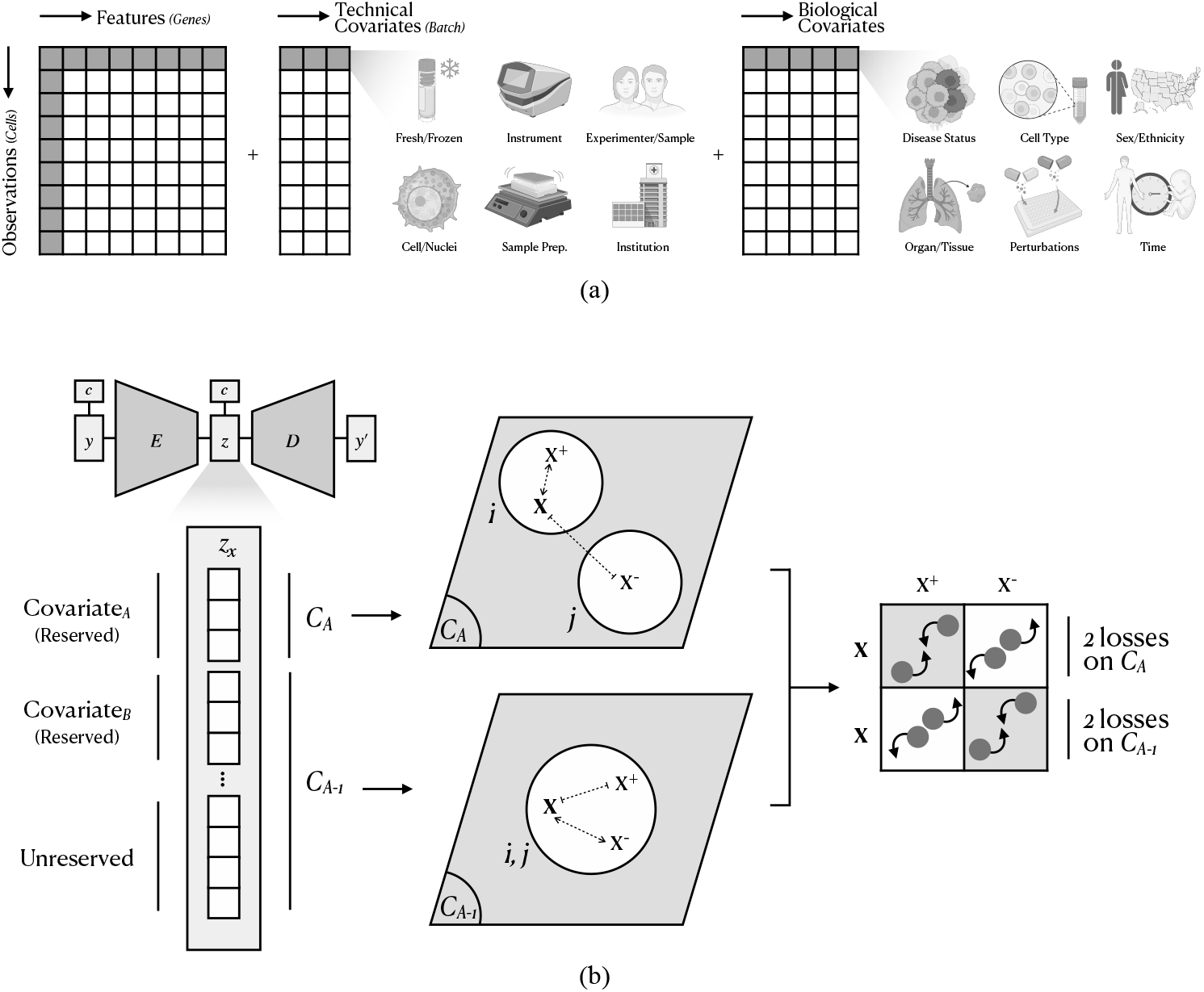
(a) A typical single-cell dataset can contain thousands or even millions of cells (observations), each with measured gene-expression profiles (features). Alongside these high-dimensional measurements, each cell is characterized by a host of technical covariates (e.g., instrumentation or sample preparation) and biological covariates (e.g., disease status, cell type, sex, ethnicity, organ, perturbations, and temporal factors). Collectively, these covariates influence gene expression, making their accurate representation and disentanglement crucial for extracting meaningful biological in-sights from complex single-cell datasets. (b) A conceptual illustration of our proposed method. The figure depicts data containing multiple covariates, denoted as *A, B*, and so on. Covariate *A* has two categories, *i* and *j*, and the current training iteration uses a data point *x* associated with category *i*. For example, Covariate *A* may represent the disease status of cells, with *i* corresponding to diseased cells and *j* to healthy cells. Here, *x*^+^ is a randomly chosen data point in category *i*, while *x*^*−*^ is a randomly chosen data point in categories other than *i*. The *C*_*A*_ (short for Covariate_*A*_, reserved latent subset of *A*) is designated to capture covariate*A*-specific information, whereas its complement *C*_*A−* 1_ (short for Covariate _*A−*1_, remaining part of the latent space) encodes all remaining variation, including other covariates and unexplained factors. The objective of the covariate-specific loss component of *A*, 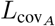, is to remove *A*-specific information from *C*_*A−*1_ and to confine only in *C*_*A*_. 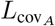 is composed of four loss terms: two operate on *C*_*A*_ to pull data points of the same category closer and push different categories apart, and two operate on *C*_*A−* 1_, which acts opposite of the losses on *C*_*A*_. By constructing and applying such sets of losses for each covariate, *TarDis* produces distinct latent subsets dedicated to individual covariates, alongside a separate space for residual variation.

Hence, it becomes essential to disentangle invariant and spurious correlations for single-cell data integration, where spurious correlations often obscure biological signals. The disentanglement of these elements not only enhances data integration by clarifying underlying biological processes but also bolsters out-of-distribution (OOD) prediction capabilities [4, 54]. Furthermore, there is a compelling need for researchers to explore the potential effects of one covariate on another, whether categorical or continuous, by manipulating such disentangled latent representations. For instance, adjusting the continuous *‘drug dose’* representation while holding the representations of *‘disease state’, ‘patient’*, and continuous *‘age’* constant could reveal the dose-dependent effects on gene expression independent of the disease’s progression or patient characteristics. Such analyses would deepen our understanding of the interactions between various factors at the cellular level, thereby unlocking new avenues for complex, hypothesis-driven research with single-cell genomics data.

To address the complexities inherent in multi-domain and multi-condition datasets, we introduce *TarDis*, a novel end-to-end deep generative model specifically designed for the *tar*geted *dis*entanglement of multiple covariates, such as those encountered in extensive single-cell genomics data. ^1^ *TarDis* employs covariate-specific loss functions through a self-supervision strategy, enabling the learning of disentangled representations that achieve accurate reconstructions and effectively preserve essential biological variations across diverse datasets. It eschews additional architectural complexities, enabling straightforward application to large datasets. *TarDis* ensures the independence of invariant signals from noise, enhancing interpretability that is crucial for extracting biological insights obscured by spurious data correlations. *TarDis* handles both categorical and, notably, continuous variables, demonstrating its adaptability to diverse data characteristics and allowing for a granular understanding and representation of underlying data dynamics within a coherent and interpretable latent space. This capability is instrumental for exploring complex biological phenomena and conducting hypothesis-driven research. Empirical benchmarking across multiple datasets highlight *TarDis*’s superior performance in covariate disentanglement, data integration, and out-of-distribution predictions, significantly outperforming existing models. ^2 3^

## 2 Results

### *2.1 TarDis* achieves robust disentanglement of covariates into isolated latent spaces

We assessed the *TarDis* model’s ability to disentangle covariates using the *Afriat* single-cell genomics dataset, which includes three distinct covariates: age, zone status, and time (Section 4.4.5). Experiments were conducted with two methodologies: disentangling all covariates simultaneously, *TarDis*_multiple_^4^, and disentangling each covariate individually followed by concatenating the reserved latent spaces, *TarDis*_single_. The disentanglement performance was benchmarked using the maximum mutual information gap (maxMIG), as detailed in Section 4.5.2, demonstrating that both configurations of *TarDis* surpassed existing models [19, 21, 57, 70, 78, 103] and achieved nearly 0.9 maxMIG scores on validation sets (Figure 2a). These results underscore the efficacy of *TarDis* in handling multiple covariates simultaneously without compromising disentanglement quality. Further analysis using the mutual information (MI) metric reveals minimal differences in the preservation of information within the unreserved and reserved latent spaces between the two training strategies, indicating the model’s effective scalability for disentanglement tasks (Figure 2b). Notably, for all subsequent experiments detailed in this paper, we have exclusively employed the multiple-covariate disentanglement approach.

**Figure 2.**
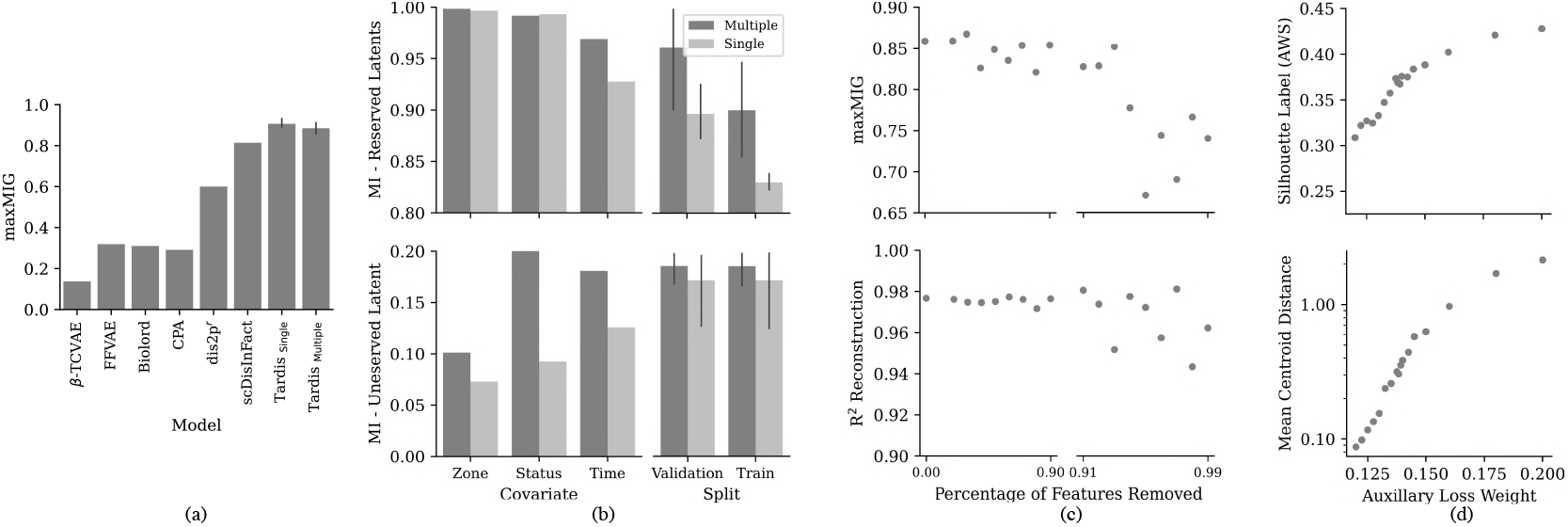
(a) Comparison of disentanglement performance using maximum mutual information gap (maxMIG), as detailed in Section 4.5.2, showing that both *TarDis* variants outperform existing models, where *r* stands for the *reported* score by the authors. Experiments (in Figure panels a and b) assessed *TarDis*_multiple_, disentangling all covariates simultaneously, and *TarDis*_single_, disentangling individually and concatenating latent spaces, with *TarDis*_single_ performing marginally superior. (b) MI in the reserved, **z**_*nk*_, and unreserved, **z**_*n*0_, latent spaces for *TarDis* under multiple and single covariate training conditions across various covariates and data splits. (c) Relationship between the percentage of input features removed and the corresponding maxMIG and R^2^ reconstruction scores, indicating robustness to feature removal. (d) Impact of auxiliary loss weight (*λ*_C_) on mean centroid distance in reserved latents, **z**_*nk*_, and average silhouette width (ASW) scores at the unreserved latent, **z**_*n*0_.

An ablation study was performed to evaluate the model’s robustness against feature reduction, where varying percentages of input features were systematically removed. Results in Figure 2c show that *TarDis* maintained high maxMIG and R^2^ reconstruction scores, above 0.65 and 0.94 respectively, affirming its resilience to input variability. Additionally, modifying the auxiliary loss weight, *λ*_C_, systematically influenced the clustering quality and disentanglement accuracy, as indicated by the increased maxMIG score and mean centroid distance with higher *λ*_C_ values (Figure 2d and Supplementary Figure 8). Moreover, the silhouette scores, calculated on the unreserved latent space **z**_*n*0_ using cell type annotations as the labels, provided empirical evidence that effective disentanglement correlates with enhanced biological signal representation, as further investigated in Results 2.2. Overall, these results not only validate the robustness of *TarDis* in disentangling complex covariate structures but also highlight its utility in preserving essential biological variations, pivotal for advancing single-cell genomic data analysis.

### 2.2 *TarDis* achieves superior performance in single-cell genomics data integration

To probe the efficacy of invariant representation learning, we turned our attention to the *Suo* dataset, a massive single-cell genomics dataset capturing human embryonic development. This dataset includes about 850k cells from various organs and time points, using multiple methods, instruments, samples, and platforms, as well as a wide range of cell types (Section 4.4.5). Its complexity makes it an ideal testbed for evaluating model performance in integrating intricate datasets. We assessed the data integration quality using the scIB package metrics [60], which are recognized benchmarks in the single-cell genomics community for evaluating the balance between biological signal preservation and batch effect mitigation (Section 4.5.2). This balance is crucial as inadequate correction can lead to data clustering by batch, obscuring true biological variance, while over-correction may suppress biological signals, reducing the biological relevance of the outcomes.

In our experimental setup, we tested two configurations of the *TarDis* model. *TarDis*-1 focuses on covariates typically considered as batch keys in single-cell data integration tasks, such as library platform, donor, sample status, and instruments. *TarDis*-2 extends this disentanglement to additional covariates including sex, age, and notably, organ. The comparative results, detailed in Table 1, show that *TarDis*, particularly *TarDis*-2, outperforms state-of-the-art models^5^ and maintains an optimal balance between biological conservation and batch correction. By effectively disentangling various spurious correlations from invariant biological signals, *TarDis* has demonstrated its robust capability to manage the complexities inherent in vast and heterogeneous datasets.

**Table 1:** Benchmarking data integration performance by scIB package [60] metrics, organized into biological signal conservation and batch correction categories (Section 4.5.2). Quantification employed a comprehensive set of metrics, with aggregate scores derived according to scIB standards. Cell-type annotations are incorporated in the metrics where *labels* are necessary. Available technical covariates—such as library platform, donor, sample status, and instrument—are concatenated and used as *batch keys* for model training, except for PCA. For this experiment, two configurations of the *TarDis* model were tested: *TarDis*-1 targets the *batch keys* defined in this experiment, while *TarDis*-2 expands disentanglement to also include sex, age, and organ. For both *TarDis* variants, the unreserved latent subsets are used for benchmarking with scIB metrics.

|  | Metric | PCA | Harmony | scVI | scANVI | inVAE | *TarDis-1* | *TarDis-2* |
| --- | --- | --- | --- | --- | --- | --- | --- | --- |
| Bio conservation | Isolated Labels | 0.610 | 0.563 | 0.638 | 0.774 | 0.798 | 0.662 | 0.767 |
|  | K-means NMI | 0.691 | 0.620 | 0.649 | 0.792 | 0.651 | 0.634 | 0.713 |
|  | K-means ARI | 0.226 | 0.182 | 0.209 | 0.360 | 0.191 | 0.185 | 0.228 |
|  | Silhouette Label (AWS) | 0.504 | 0.482 | 0.496 | 0.576 | 0.508 | 0.497 | 0.508 |
|  | Cell-type LISI (cLISI) | 0.999 | 0.997 | 0.999 | 1.000 | 0.999 | 0.998 | 0.999 |
| Batch correction | Silhouette Batch | 0.851 | 0.862 | 0.867 | 0.861 | 0.840 | 0.903 | 0.896 |
|  | Integration LISI (iLISI) | 0.057 | 0.100 | 0.098 | 0.093 | 0.040 | 0.094 | 0.080 |
|  | kBET per Label | 0.309 | 0.475 | 0.487 | 0.526 | 0.194 | 0.448 | 0.430 |
|  | Graph Connectivity | 0.793 | 0.671 | 0.866 | 0.912 | 0.836 | 0.866 | 0.879 |
|  | PCR Comparison | 0.000 | 0.350 | 0.699 | 0.222 | 0.000 | 0.931 | 0.850 |
| Aggregate score | Bio conservation | 0.606 | 0.569 | 0.598 | **0.701** | 0.629 | 0.595 | 0.643 |
|  | Batch correction | 0.402 | 0.492 | 0.603 | 0.523 | 0.382 | **0.648** | 0.627 |
|  | Total | 0.524 | 0.538 | 0.600 | 0.629 | 0.530 | 0.616 | **0.637** |

### *2.3 TarDis* generates ordered latent representation for continuous covariates

In addressing the challenge of learning the representation of disentangled *continuous* covariates, *TarDis* provides a solution that captures data variations without reducing them to mere categorical approximations. Continuous covariates such as age or treatment dosage are critical for understanding gradients in biological processes, cellular behavior, and disease progression. To manage the subtleties associated with these variables, *TarDis* employs a distance-based loss function for each auxiliary loss component. The model employs negative pair losses weighted by the distance between the values of the continuous covariates, omitting positive pair losses due to the continuous nature of the covariate, which results in generating an ordered and interpretable latent space (Figure 3).

**Figure 3.**
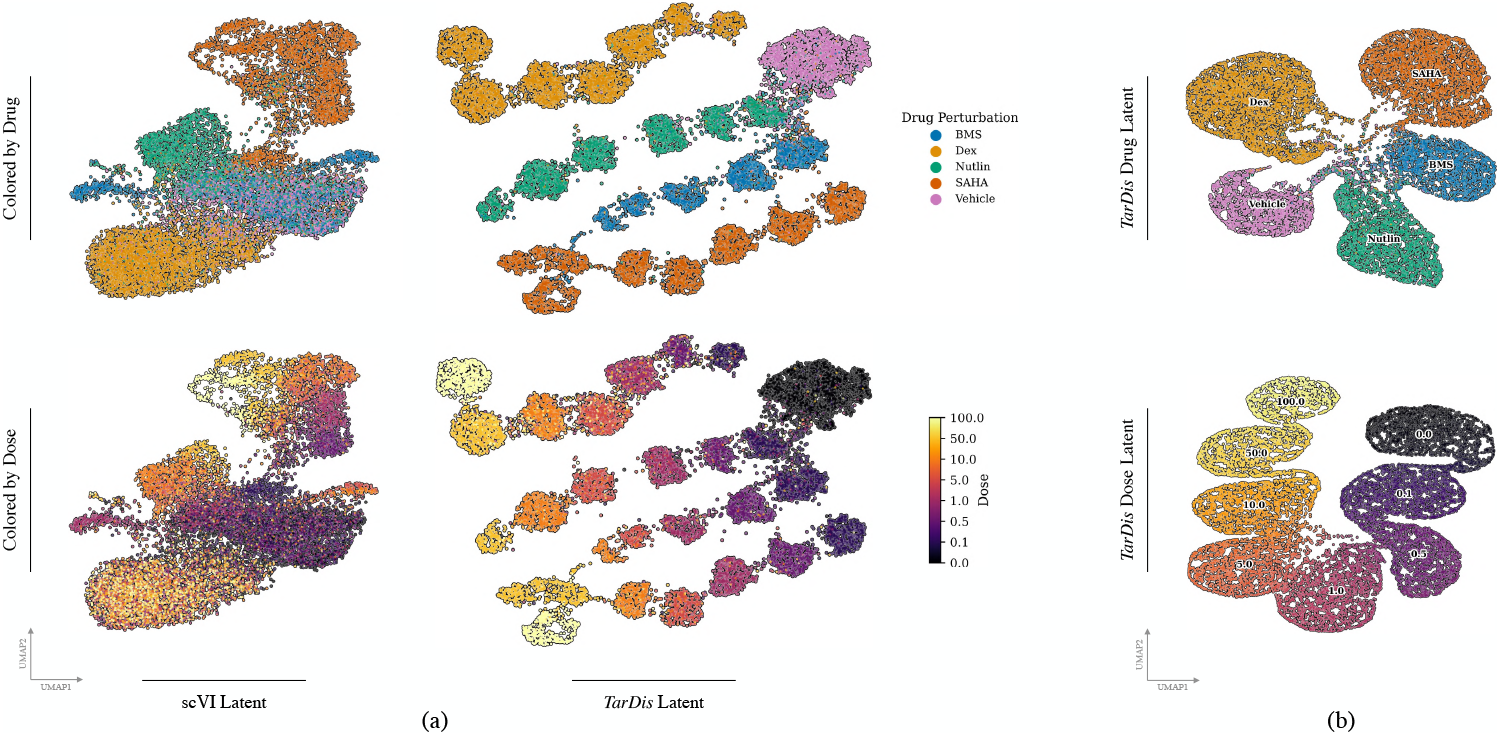
UMAP visualization [62] of *TarDis* latent space representations from the *Sciplex* dataset (a) Comparing the performance of scVI (trained using default parameters from the scvi_tools package [25], with batch keys included) and *TarDis* models in capturing drug responses and dosage effects. The upper row displays clusters differentiated by drug types, while the bottom row illustrates the ordered representation of dosage, showcasing the ability of *TarDis* to structurally organize cellular responses across different drug concentrations. (b) *TarDis* model training generates three distinct latent spaces: unreserved, dose, and drug. Displayed UMAPs are the dose and drug latent subspaces, demonstrating structured separation and ordered representation.

In our studies, we focused on two primary continuous covariates, age and drug dosage, which present distinct challenges due to their variability and significant impact on cellular phenotypes. We employed two datasets to evaluate the effectiveness of *TarDis* in producing ordered latent representations of these covariates. The first dataset, named *Sciplex* (Section 4.4.5), involves drug perturbation experiments and helps in analyzing the structured response of cells to varying drug dosages. The second dataset, referred to as *Braun* (Section 4.4.5), comprises 1.6 million cells from human embryonic brain development, providing a complex scenario for assessing the impact of time as a continuous variable. Through *TarDis*, we managed to produce ordered latent representations of these covariates within isolated latent subsets while concurrently disentangling other variables such as the type of library platform, donor characteristics, sample status, instrumentation used, and tissue types (Figure 3, 4).

**Figure 4.**
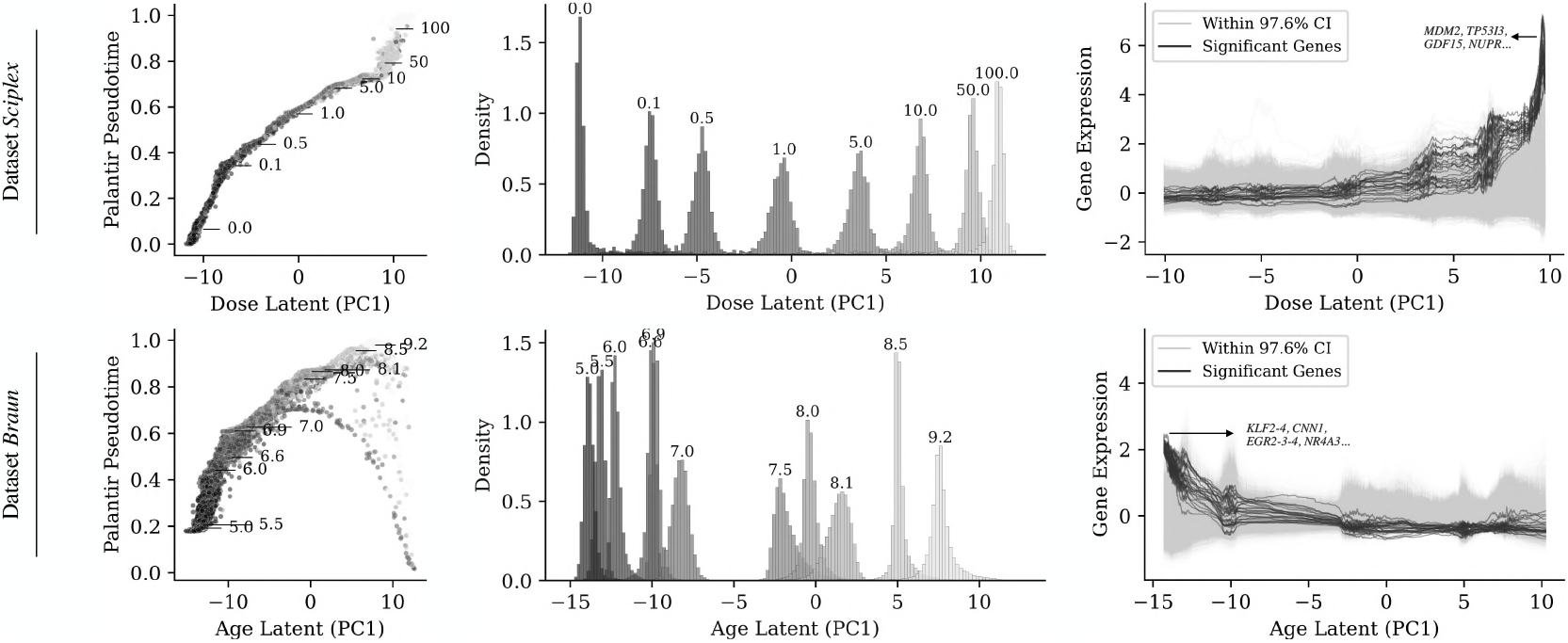
Ordered latent spaces for dose and age (post-conception week) in the *Sciplex* and *Braun* datasets, respectively. (left) Principal Component 1 (PC1) of the continuous covariate latent space plotted against Palantir pseudotime [77], which uses a k-nearest neighbor graph to infer cell pseudotime trajectories. (middle) Density distribution of the continuous covariate in the respective latent subset, illustrating ordered peaks corresponding to varying levels of the covariate. (Right) Differential gene expression profiles plotted against the continuous covariate latent space, identifying genes that show variation in expression levels associated with changes in the covariate, indicative of underlying cellular processes. Gene expression patterns are highlighted with (upper right) increasing doses of *Nutlin* and (bottom right) through human embryonic developmental stages of *forebrain neuron*.^6 7^

This representation has enabled previously unfeasible hypothesis-driven biological analyses. For example, *TarDis* allows for the exploration of organ-specific developmental gene expression patterns for specific cell types, an analysis that previously wasn’t optimal with non-batch-corrected input spaces. Unlike existing models such as scVI and scANVI, which address batch effects but often fail to retain essential biological information like age or organ specifics —either being overly corrected by batch keys or inadequately accounted for [55, 97]— *TarDis* allows researchers to isolate cells from two different organs using the organ-specific latent subset and, for a given cell type, compare expression patterns across developmental stages in a massive multi-organ developmental single-cell dataset. This analysis benefits from a batch-corrected latent space, thanks to a set of other latent subsets that disentangle batch effects ^6^. In Figure 4 bottom right, *TarDis* enabled to identify genes including `EGR2-3-4`, `KLF2-4`, `RTL1`, `SPRY4-AS1`, and `FOSB`, that decrease in expression through embryonic development of human *forebrain neurons* within the *Braun* dataset, which were shown to be associated with brain development, aging, and diseases including Down syndrome and bipolar disorder [20, 46, 61, 68, 71, 98]. In a parallel experiment using the *Sciplex* perturbation dataset, *TarDis* effectively disentangled the influences of drug type and dosage (Figure 3, 4). Using the data points corresponding to *Nutlin* cluster in drug latent, we analyzed how gene expression responds to increasing doses. As shown in Figure 4 upper right, this approach allowed us to pinpoint the expression patterns of genes such as `TP53I3`, `CDKN1A`, `GDF15`, `MDM2`, `FDXR`, and `NUPR1`, which are notably responsive to escalating doses of *Nutlin* [36, 93]. ^7^ ^8^

### *2.4 TarDis* predicts counterfactual gene expressions accurately under OOD conditions

The capacity of predictive models to generate accurate gene expressions under OOD conditions is pivotal for extrapolating research findings to new or novel environments. In evaluating this capacity, *TarDis* was systematically tested using two distinct datasets to gauge its effectiveness in predicting counterfactual gene expressions. Using the *Afriat* dataset, previously introduced, multiple models were trained, each excluding a different combination of three covariates to create respective OOD sets. Additionally, the *Miller* dataset, which comprises samples from human developmental embryonic lung, was utilized to disentangle the effects of age and donor covariates (Section 4.4.5). Similar to the *Afriat* dataset, combinations of two covariates were systematically omitted during training to simulate various OOD conditions.

*TarDis* demonstrated superior performance in predicting gene expressions under OOD conditions, outperforming CPA^5^, another model that *concurrently* disentangles multiple covariates [57], in both the *Afriat* and *Miller* datasets. In the *Afriat* dataset, *TarDis* achieved notably higher R^2^ reconstruction scores, showcasing its strong capability for accurate reconstruction under varied and unseen conditions (Section 4.5.2). In the *Miller* dataset, the challenge intensified with the evaluation focusing on differentially expressed genes, DEGs (Section 4.5.2). *TarDis* excelled, achieving significantly better OOD predictions for DEGs compared to CPA. Notably, our inclusion of both R^2^ and R^2^-DEG provides a dual-resolution view, where broad data distribution reconstructions and global patterns across the entire gene set are highly accurate. Simultaneously, the subset of genes undergoing biologically meaningful expression shifts remain well-captured, ensuring that critical biological signals crucial for downstream analyses are preserved. These results, shown in Figure 5, affirm the utility of *TarDis* not only in disentangling complex covariate interactions within datasets but also in its capability to generalize across novel, unseen domains, key to advancing the precision and reliability of predictive models in single-cell genomics.

**Figure 5.**
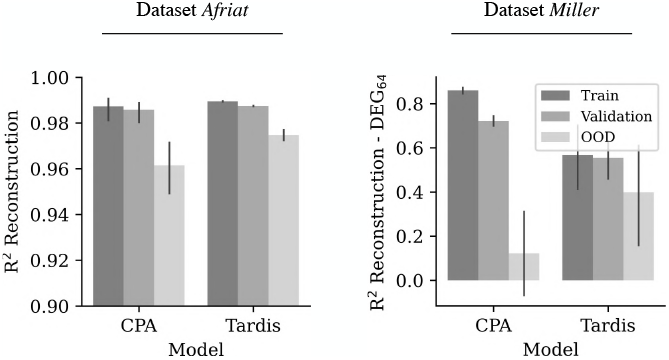
Performance comparison of *TarDis* and CPA in predicting counterfactual gene expressions under out-of-distribution conditions using the *Afriat* and *Miller* datasets. R^2^ scores for reconstructed gene expressions and differentially expressed genes (DEG) across varying unseen covariate combinations highlight *TarDis*’s superior predictive capabilities.

### *2.5 TarDis* produces interpretable latent representations of disentangled covariates

In exploring the capabilities of *TarDis* to yield interpretable latent representations, we utilized the *Norman* dataset, a comprehensive collection comprising 108k cells subjected to single or combinatorial gene perturbations (Section 4.4.5). This dataset is particularly challenging due to the diversity and complexity of its perturbations, with a total of 284 distinct perturbation conditions included in this analysis. In this experiment, the inference model in *TarDis* relied solely on input features without the introduction of covariate information, **s**_*n*_. This approach ensured that the learning process was purely driven by the data’s inherent structure rather than external annotations. Our results indicate that *TarDis* effectively disentangles these perturbations, with each perturbation distinctly isolated in the latent space. Significantly, perturbations that share a common cellular program, as identified in the original publication of the dataset [66], were found to cluster closely. The results support *TarDis* ability to capture interpretable and biologically meaningful patterns, as the clustering is not random qualitatively but reflects the underlying biological relationships (Figure 6a).

**Figure 6.**
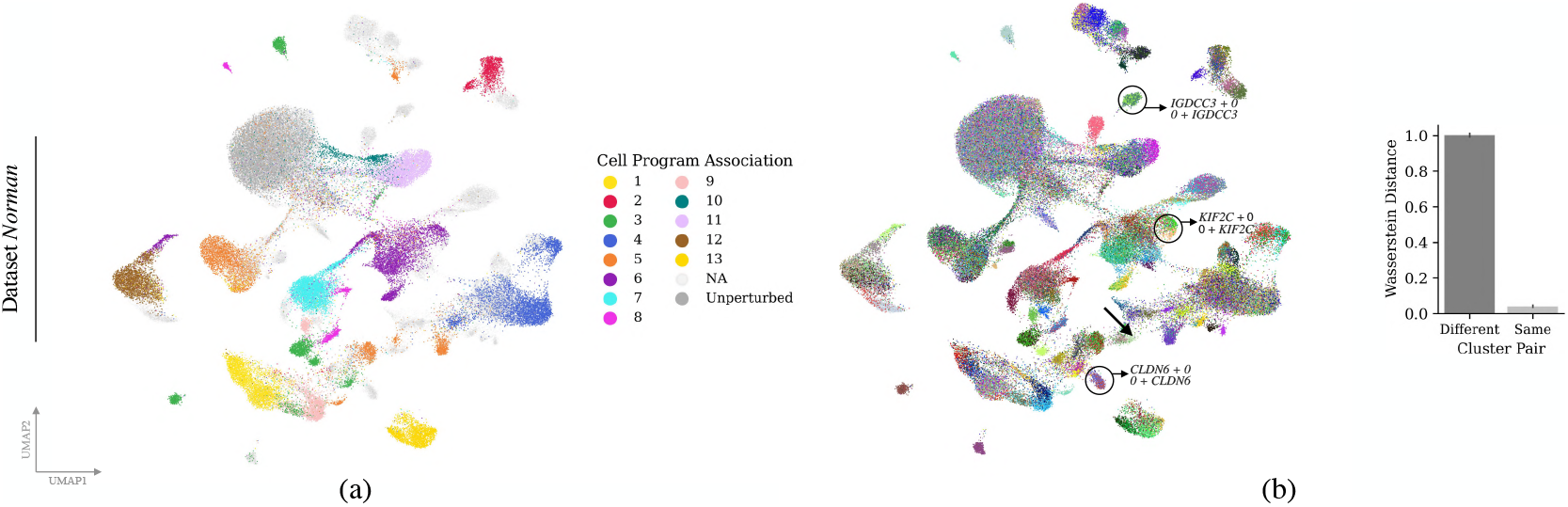
UMAP visualization of the *TarDis* perturbation latent space derived from the *Norman* dataset: (a) Clusters corresponding to sets of perturbations associated with similar cell programs as identified in the original publication [66], demonstrating the model’s ability to capture underlying biological patterns. (b) UMAP visualization of *TarDis* latent space, colored by 270 perturbations. Representative clusters are highlighted, illustrating the model’s capability to align identical perturbations accurately despite nominal labeling differences, thus confirming label reconciliation. Wasserstein distances are computed to quantitatively confirm the close, often overlapping, clustering of identical perturbations [90].

A particularly rigorous test of the model’s interpretability involved the re-labeling of certain perturbations in the dataset. Specifically, the labels were altered to appear as two distinct entities: *‘X+0’* and *‘0+X’*, despite originating from the same perturbation. This was designed to test whether *TarDis* could recognize and reconcile these as identical despite their nominal differences. The results were in line with our expectations: *TarDis* successfully overlapped these perturbations in the latent space, affirming its capability to generate biologically coherent and interpretable latent representations, even under challenging conditions (Figure 6b). This analysis not only confirms the robustness of *TarDis*’s disentanglement capabilities but also highlights its potential in generating actionable insights from complex genomic data, where interpretability is crucial for meaningful biological inference.

## 3 Conclusion

In this study, we presented *TarDis*, a novel deep generative model designed for the targeted disentanglement of covariates in complex multi-domain and multi-condition datasets, particularly focusing on the challenges presented by single-cell genomics data. Our approach leverages a series of covariate-specific loss functions to facilitate robust disentanglement and invariant representation of both continuous and categorical variables, thus enhancing data integration capabilities and enabling more insightful biological analyses. Through rigorous benchmarking against existing models and diverse datasets, *TarDis* has demonstrated superior performance not only in its capacity to disentangle complex covariate structures but also in maintaining essential biological signals crucial for accurate data interpretation and analysis, and generating robust predictions under out-of-distribution conditions. Moreover, *TarDis*’s ability to generate ordered latent representations of continuous covariates significantly enhances differential analyses across varying conditions. The model perform robustly in generating interpretable and biologically meaningful latent representations, which could empower researchers to conduct advanced hypothesis-driven research, potentially unveiling novel insights and therapeutic targets.

Including all covariates may appear simpler *a priori*, but it often induces conflation of correlated factors, leakage into the unreserved latent space, and diminished performance and interpretability (Figures 2 and ‘Limitations’ under ‘Methods’ section). Consequently, *hypothesis-driven selection* of a subset of covariates—guided by the overarching research question—remains the most direct and scientifically productive strategy for employing *TarDis*. By aligning the targeted covariates with biological or clinical priorities, researchers can extract clearer insights, avoid overcorrection, and preserve critical aspects of the data that remain unreserved for other analyses.

*TarDis* establishes a robust approach for exploring complex biological questions, offering researchers unprecedented clarity in dissecting the nuanced interactions between diverse covariates. This capability is instrumental in advancing personalized medicine, supporting the development of customized therapeutic strategies grounded in a profound understanding of individual responses to different treatments. Considering the expansion of *TarDis* applications beyond genomics, for instance into neuromarketing using EEG event-related potentials (ERP) data, it becomes crucial to acknowledge that modifications to the model may be necessary to accommodate different types of data. We are actively investigating these potential applications, aiming to extend the reach and impact of *TarDis* across various scientific and applied fields^9^.

## 4 STAR Methods

### 4.1 Key resources table

**Table 2.**

| Reagent or Resource | Source | Identifier |
| --- | --- | --- |
| **Deposited data**<sup>10</sup> |  |  |
| *Afriat dataset*, single-cell RNA sequencing data of host and parasite gene expression during the liver stage of the rodent malaria parasite *Plasmodium berghei* ANKA. | Afriat et al., 2022 [1] | GSE181725 (GEO) |
| *Suo dataset*, a multi-organ, single-cell transcriptomic perspective, capturing dynamic immune system developments across nine prenatal human tissues during embryonic stages | Suo et al., 2022 [86] | E-MTAB-11343 (ArrayExpress) |
| *Braun dataset*, a comprehensive single-cell transcriptomic analysis of the human brain during the crucial first trimester | Braun et al., 2023 [12] | EGAS00001004107 (European Genome Phenome Archive) |
| *Miller dataset*, provides a detailed single-cell mRNA sequencing atlas of human lung development from 11.5 to 21 weeks, integrated with studies on homogeneous human bud tip organoid cultures | Miller et al., 2020 [63] | E-MTAB-8221 (ArrayExpress) |
| *Sciplex dataset*, derived from the sci-Plex technology using nuclear hashing, quantifies transcriptional responses to chemical perturbations at single-cell resolution | Srivatsan et al., 2020 [84] | GSE139944 (GEO) |
| *Norman dataset*, leverages high-content Perturb-seq technology to explore cellular and organismal complexity through combinatorial gene expression | Norman et al., 2019 [66] | GSE133344 (GEO) |
| **Software and algorithms** |  |  |
| TarDis | This study | theislab/tardis (GitHub) |
| β-TCVAE | Chen et al., 2018 [19] | rtqichen/beta-tcvae (GitHub) |
| FFVAE | Creager et al., 2019 [21] | nomnomnonono/FFVAE (GitHub) |
| Biolord | Piran et al., 2024 [70] | nitzanlab/biolord (GitHub) |
| CPA | Lotfollahi et al., 2023 [56] | theislab/cpa (GitHub) |
| scIB | Luecken et al., 2022 [60] | YosefLab/scib-metrics (GitHub) |
| scVI | Lopez et al., 2018 [55] | scverse/scvi-tools (GitHub) |
| scANVI | Xu et al., 2021 [97] | scverse/scvi-tools (GitHub) |
| Harmony | Korsunsky et al., 2019 [49] | lilab-bcb/harmony-pytorch (GitHub) |
| scDisInFact | Zhang et al., 2024 [103] | ZhangLabGT/scDisInFact (GitHub) |
| inVAE | Aliee et al., 2023 [4] | theislab/inVAE (GitHub) |
| **Other** |  |  |

<sup>10</sup>Refer to Section 4.4.5 for a detailed description of the datasets used and their characteristics.

Table 2 continued from previous page
|  |  |  |
| --- | --- | --- |
| Mamba environment for reproducibility, including all used software and their versions | This study | [theislab/tardis/environment](https://github.com/theislab/tardis/environment) (GitHub) |

### 4.2 Resource availability

#### 4.2.1 Lead contact

Further information and requests for resources should be directed to and will be fulfilled by the lead contact, Kemal Inecik.

#### 4.2.2 Materials availability

This study did not generate new materials.

#### 4.2.3 Data and code availability

- This manuscript utilizes pre-existing datasets that are publicly accessible; the specific accession numbers pertinent to these datasets are provided in the key resources table in Section 4.1. Detailed descriptions of the dataset contents, along with alternative sources for data acquisition, are presented in Section 4.4.5. Comprehensive documentation of the preprocessing procedures applied to the majority of these datasets is available in the GitHub repository of *TarDis*.
- All original code has been deposited at GitHub repository (theislab/tardis) and is publicly available as of the date of publication.
- For any supplementary information necessary to execute the model training processes of *TarDis*, interested parties are encouraged to contact the lead author, who will provide the required details upon request..

#### 4.2.4 Resource availability

This paper analyzes existing, publicly available data. These accession numbers for the datasets are listed in the key resources table in Section 4.1.

### 4.3 Method details

#### 4.3.1 Overview

Let 𝒟 represent a single-cell genomics dataset containing *N*_*C*_ cells, where each cell, denoted as *n*, is characterized by its gene expression (**x**_*n*_) and associated covariates (**s**_*n*_). The gene expression is represented by a count vector 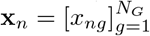, where *x*_*ng*_*∈* ℤ_*≥*0_ is the expression count of gene *g*, and *N*_*G*_ is the total number of genes in the dataset. Additionally, each cell *n* is associated with a vector of covariates 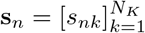, which may be either continuous or discrete, and *N*_*K*_ indicates the number of covariates. *TarDis* constructs a latent representation **z**_*n*_ for gene expression **x**_*n*_, organized as 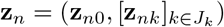, where *J*_*k*_ ⊆ {1, …, *N*_*K*_} denotes the subset of covariates targeted for disentanglement. Note that since indexing of targeted covariates uses *k ∈ J*_*k*_ starting from 1, the subscript ‘0’ in **z**_*n*0_ does not correspond to any index in *J*_*k*_ but instead represents the portion of the latent space reserved for information not explained by the targeted covariates. Specifically, **z**_*nk*_ is a latent vector constructed for each targeted covariate, while **z**_*n*0_ captures residual information independent of targeted covariates. During model training, *TarDis* employs a novel approach to foster disentanglement by generating pairs of additional latent vectors 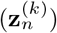 and 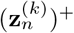 corresponding. to two data points 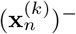 and 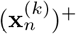 These data points are selected *randomly* to two data points and differ in the *k*_th_ covariate value, such that 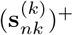 and 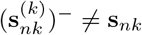. Observe that, for each categorical covariate *k*, a *positive* pair for covariate *k* consists of the original data point and another randomly selected data point from the dataset whose category matches exactly for covariate *k*; formally, 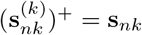. On the other hand, a *negative* pair for covariate *k* comprises the original data point and another randomly selected data point whose category differs (i.e., belongs to any category other than the original) for covariate *k*. Formally, 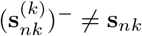. These definitions are consistently maintained in all subsequent formulations involving pair-wise terms in the loss function.

The primary objective of *TarDis* training is to optimize the latent vectors based on a distance measure *F*. While *F* is defined conceptually as a real-valued function, 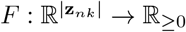, here just to illustrate the underlying concept, practical implementation typically employ multiple loss terms instead of a single function for optimizing latent vectors, as will be discussed in further detail. For each covariate *k ∈ J*_*k*_, *F* should satisfy 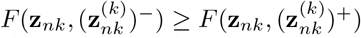, implying that latent vector **z**_*nk*_ should be more similar to another vector that shares the same covariate value, 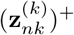 than to a vector with a different covariate value, 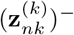. Furthermore, the latent vector **z**_*n*0_ should show equal similarity to any other vectors regardless of their covariate values, whether 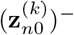 and 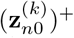, thus fulfilling the condition: 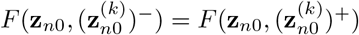 This equality ensures that **z**_*n*0_ remains unaffected by covariate-specific information, thereby providing a covariate-neutral representation of the cell’s gene expression. Ultimately, the aim of *TarDis* is to produce a latent representation in which **z**_*nk*_ reflects the influence of its corresponding covariate **s**_*nk*_, while **z**_*n*0_ offers a covariate-neutral representation of the cell’s gene expression profile, unaffected by any covariate-specific variations ^11^.

#### 4.3.2 VAE skeleton

*TarDis* builds upon a variational autoencoder (VAE) to construct a high-fidelity generative model that underpins our disentanglement objectives. The VAE component optimization guided by the Evidence Lower Bound (ELBO), a surrogate for the intractable marginal log-likelihood as shown in Equation 1 [45]. Here, the covariates, **s**_*n*_, are pivotal for capturing factors that might influence the observed data, such as batch effects. *TarDis* incorporates the target covariates as **s**_*n*_, and also allows inclusion of non-target covariates, providing flexibility in managing different types of data impacts. The first term of ℒ_VAE_ represents the reconstruction loss, ℒ_R_, which quantifies the expected negative log-likelihood of the observed data **x**_*n*_, conditioned on the latent variables, **z**_*n*_. It aims to minimize the discrepancy between the observed data and its reconstruction from the latent space. The reconstruction loss is formally expressed using the negative binomial (NB) distribution, ideal for capturing the count variability inherent in data types like single-cell genomics (Equation 2). In this equation, Γ denotes the gamma function, ***µ*** and ***θ*** refer to the mean and inverse dispersion parameters of the negative binomial distribution, respectively [40]. The second term measures the Kullback-Leibler divergence (KL), ℒ_KL_, penalizing deviations of the learned posterior distribution *q*_*ϕ*_(**z**_*n*_ | **x**_*n*_, **s**_*n*_) from the prior distribution *p*(**z**_*n*_). In Equation 3, the approximate posterior distribution is assumed to be Gaussian distribution with mean ***µ***_*n*_ and diagonal covariance matrix **Σ**_*n*_, and the prior distribution *p*(**z**_*n*_) is typically a standard normal distribution 𝒩(**0, *I***) where ***I*** is the identity matrix in 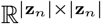. This acts as a regularizer, aligning the latent embeddings with a predefined distribution, to ensure the statistical robustness and generalization capability of the model [45].

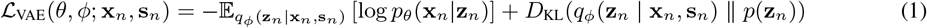

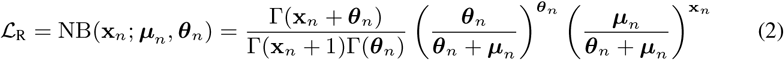

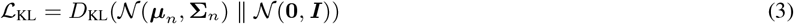

#### 4.3.3 Auxiliary loss

In *TarDis* model training, the VAE optimization is intertwined with the novel auxiliary loss component introduced, ℒ_C_, to construct 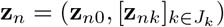 with **z**_*nk*_ ~ 𝒩(***µ***_*nk*_, **Σ**_*nk*_). The overall loss function of *TarDis* integrates these components through a weighted sum, controlled by hyperparameters *λ*_C_, *λ*_KL_, and *λ*_R_ (Equation 8). Specifically, ℒ_C_ is a composite loss function that incorporates four distinct loss components for each covariate[13]. For each target covariate **s**_*nk*_, the loss function, 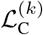, includes 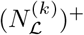 positive and 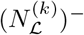 negative loss terms. Similarly, for the covariate-free representation **z**_*n*0_, it includes 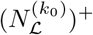 positive and 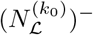 negative terms. The losses for positive pairs and negative pairs given in Equations 4 and 5, respectively. Here, the *λ* values are hyperparameters that determine the weight of each loss component, while the ℒ loss functions encompass metrics such as KL divergence and mean squared error (MSE)^12^. Thus, the overall covariate loss, 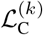, is computed as the sum of these two pair losses, as specified in Equation 6. By aggregating these individual covariate losses, the total auxiliary loss, ℒ_C_, is expressed in Equation 7.

The configuration of 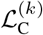 is meticulously designed to meet several critical objectives within the *TarDis* framework. First, by minimizing the distance between 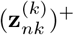 and **z**_*nk*_, the model ensures that the latent representations of positive examples closely align with their corresponding covariate within respective latent subset, accurately reflecting specific characteristics. In contrast, it maximizes the distance between 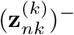 and **z**_*nk*_, thereby promoting clear differentiation in the latent representations of negative examples and enhancing the distinction between different covariates. Additionally, the model strategy involves maximizing the distance between 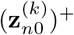 and **z**_*n*0_, while minimizing the distance between 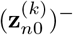 and **z**_*n*0_. This approach ensures that **z**_*n*0_ remains free from covariate-specific influences, maintaining its role as a covariate-neutral representation. These operations collectively ensure that covariate information is precisely captured in the respective targeted latent subsets, **z**_*nk*_, and effectively isolated from **z**_*n*0_.

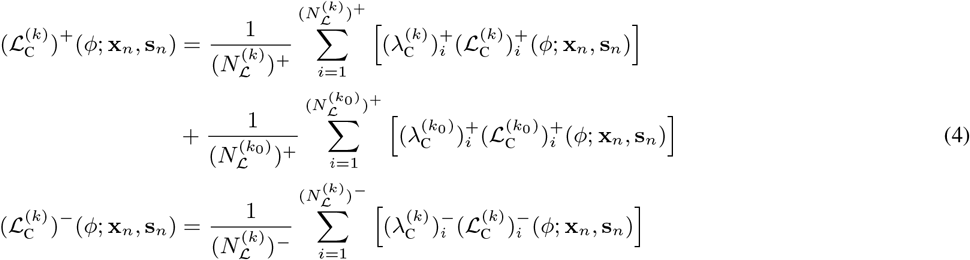

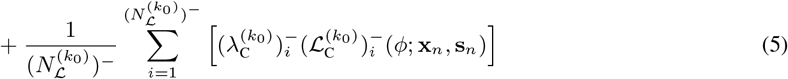

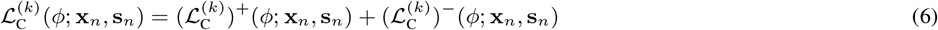

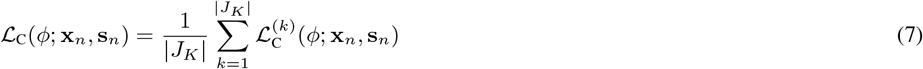

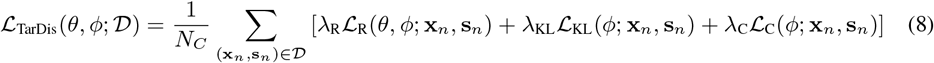

To clarify the construction of the observation-specific loss, note that the complete objective is built hierarchically by aggregating per-observation contributions across data points and covariates. Each data point (**x**_*n*_, **s**_*n*_) generates partial losses for each targeted covariate *k*, comprising a negative-pair term 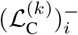 (Equation 5) and a positive-pair term 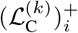 (Equation 4), where *i* indexes the respective pairs. For covariate *k*, the negative-pair loss in Equation 5 sums over 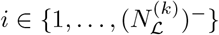 to aggregate contributions from all negative pairs, while the positive-pair loss in Equation 4 analogously sums over 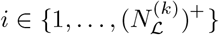. These covariate-specific components are combined in Equation 6 as 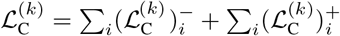, ensuring each data point’s contribution is preserved. The global auxiliary loss (Equation 7) then averages 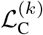 over all targeted covariates *k ∈ J*_*K*_, which is finally combined with the VAE’s reconstruction and KL terms in Equation 8 to form the total loss ℒ_TarDis_. This hierarchical formulation explicitly decomposes how individual observations, through their positive and negative pairs, influence the training objective at both covariate-specific and dataset-wide levels.

Various options for the loss functions used in 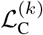 are explored. Although the theoretical framework primarily employs KL divergence as the loss metric, the principle is also applicable to various losses between anchor points and negative or positive samples with minor adjustments ^12^. Also note that, for continuous covariates, we adopt an analogous strategy but weigh the negative-pair penalty based on the distance between covariate values rather than using discrete category mismatches, while positive-pair loss components disappears. The optimization of various hyperparameters, including the individual loss weights, is conducted once and uniformly applied across all experiments, unless explicitly stated otherwise ^13^. The experiments and benchmarking processes utilize a diverse array of datasets to ensure comprehensive testing and validation of the model. Each dataset is selected to represent different types and scales of data challenges ^14^. Various evaluation metrics are used to assess the model’s performance, with a full discussion provided in Section 4.5.2 ^15^. The assumptions behind the theoretical framework are discussed in Section 4.4.4^16^.

### 4.4 Additional details

#### 4.4.1 Related Work

Models like single-cell Variational Inference (scVI) facilitate data integration by incorporating environmental variables such as experimental batches or sequencing protocols alongside gene data, using one-hot vectors processed through a conditional variational autoencoder (cVAE) to reduce technical noise [55]. Its extension, single-cell ANnotation using Variational Inference (scANVI), builds on this by introducing cell annotations in a semi-supervised approach, thus enhancing cell integration across diverse environments and adeptly capturing cell type variations [22, 97]. Despite their effective integration, these models may over-correct, adjusting biological signals while targeting technical noise, which can obscure subtle biological variations such as inter-patient differences or treatment effects. Moreover, these methodologies tend to aggregate all sources of spurious correlations indiscriminately, failing to discern the unique characteristics of each source [22, 55, 88]. This approach inadequately addresses the nuanced interactions between these sources and biological signals, particularly problematic with continuous spurious covariates such as age or drug dosage. Models equipped to continuously adapt to these subtle variations are thus essential, ensuring that biological insights derived from single-cell genomics are not confounded by these varying conditions.

Several models in single-cell genomics have explored creating multiple latent spaces to handle different sources of variability distinctly. For instance, contrastiveVI models each covariate separately, developing a shared latent space for the common variability across covariates and an exclusive latent space for the target covariate’s unique variability [95]. Similarly, single cell disentangled Integration preserving condition-specific Factors (scDisInFact) develops a shared latent space specifically designed to account for and eliminate batch effects, while simultaneously maintaining separate latent spaces for other covariates, isolating and preserving the variations from batch influences. [103]. Yet, none of these approaches offer a control latent space dedicated to retaining batch effects while filtering out the influences of other covariates, essential for accurately distinguishing between variations caused by batch effects and those arising from true biological differences. Such methods draw inspiration from broader approaches focused on fair and disentangled representation, such as Flexibly Fair VAE (FFVAE) and Fader networks, and unsupervised disentanglement techniques such as Total Correlation VAE (*β*-TCVAE) [19, 52, 67, 78]. The cell optimal transport model (CellOT) uses optimal transport (OT) methods to align cells from control and perturbed conditions, but its non-generative, single-covariate focus limits broader applicability [14]. Biolord offers a unique approach to supervised disentanglement, yet it faces scalability issues due to per-cell optimization [70]. The invariant VAE (inVAE) method introduces conditional priors within the VAE framework to effectively disentangle spurious and invariant correlations. While it offers nuanced disentanglement, inVAE faces optimization challenges, particularly in large datasets, and does not separate latent representations for individual covariates, and does not support continuous covariates naively limiting its ability to analyze complex interactions between various biological conditions in detail [4]. On the other hand, Compositional Perturbation Autoencoder (CPA) handles drug perturbations and produce latent embedding but their assumption of linearity in the latent space limits capturing complex, non-linear biological interactions [41, 56].

While existing approaches in single-cell genomics have notably advanced the disentanglement of spurious and invariant correlations, they predominantly excel within narrowly defined scenarios. Many models, however, simplify continuous covariates by categorizing them, which undermines the granularity of biological insights and limits their applicability in precision medicine. Beyond this, there’s a critical need for models that not only handle the diversity of single-cell data but also scale efficiently and train effectively given the heterogeneity inherent in these datasets. Despite the innovative nature of these methods, they are often tailored to specific experimental conditions rather than offering a universal solution across the diverse landscape of single-cell analysis. There remains an unmet need for a comprehensive model that excels in data integration, out-of-distribution prediction, and serves as a robust platform for addressing intricate biological questions across various conditions and experimental setups.

#### 4.4.2 Limitations

While *TarDis* introduces significant advancements in disentangling complex covariate structures in single-cell genomics, it is important to acknowledge several inherent limitations. *TarDis* operates under a supervised learning paradigm, which necessitates access to pre-labeled covariates. This requirement limits its applicability to datasets where such labels are readily available and accurately annotated, constraining its utility in less structured environments.

A notable limitation of *TarDis* is the potential for overfitting. Although rigorous validation protocols and robust regularization strategies, including elevated dropout rates and weight decay—more aggressive than those utilized in generic VAE models like scVI—are employed, the risk remains. In our study, the hyperparameters were carefully optimized at the onset of all experiments, ensuring consistent conditions across all tests, which mitigated the concerns of overfitting. It is important to note that our successful one-time optimization and the avoidance of overfitting in single-cell genomics data do not guarantee similar outcomes across other data types, hence users must conduct cautious benchmarking on validation splits to ensure the model’s generalizability.

Moreover, the disentanglement of interdependent covariates introduces unique challenges. For example, accurately disentangling *age* and *donor* in a single-cell genomics data as covariates requires the presence of multiple donors of varying ages to prevent the model from conflating these factors. Without such diversity, the model risks inaccurately attributing the influence of one covariate to another, thereby undermining the reliability of the disentanglement, particularly evident in our validation splits.

Additionally, the implementation of *TarDis* introduces computational overhead, slightly slowing down the processing speed. Nevertheless, this does not significantly impact performance, even with large datasets like the *Braun* dataset, which comprises 1.6 million cells. The primary bottleneck arises from the selection of counteractive minibatches for each covariate during training, which is quantified to increase the average training time by approximately 1.8 times in comparison to scVI, when three covariates were targeted.

The encoding of covariates in a one-hot format, **s**_*n*_, while optional as mentioned in Section 4, generally fosters better disentanglement in the validation splits. However, the dependency of disentanglement on the input space may necessitate further optimization. This adjustment is crucial for enhancing the model’s utility in specific downstream tasks, as demonstrated in our analysis using the *Norman* dataset in Section 2.5.

Lastly, *TarDis* necessitates numerous hyperparameters, especially concerning the loss weights for each of the four terms associated with every covariate. This complexity was manageable in our experiments through our aforementioned one-time optimization, and it did not present issues for single-cell data. However, adapting the model to new datasets could necessitate further tuning, potentially complicating its application across varied contexts. It is also important to underscore the model assumptions in Section 4.4.4, as these foundational assumptions highlight potential limitations and areas where *TarDis* might encounter challenges.

#### 4.4.3 Model training details

##### Compute Resources and System Configuration

For the computational tasks in our research, we employed *NVIDIA Tesla A100* GPUs, which feature 40 GB of high-bandwidth HBM2 memory each. This GPU architecture is specifically designed for accelerating machine learning and high-performance computing applications, providing substantial throughput for both single and mixed-precision computations. We allocated 64 GB of GPU memory for processing large training datasets, which facilitated efficient handling of extensive computational operations without the need for frequent data swapping, thereby minimizing I/O overhead. For smaller datasets, a reduced memory allocation of 16 GB was used, which optimized resource utilization without compromising performance. On the CPU side, our computational nodes were equipped with dual *Intel Xeon Gold 6230* processors. Each processor offers 20 cores operating at a base frequency of 2.1 GHz, which can boost up to 3.9 GHz. This setup provided a robust and responsive environment for handling non-GPU-intensive tasks and managing the preprocessing and postprocessing stages of our experiments. The system’s main memory configuration included 256 GB of DDR4 RAM per node, which was crucial for supporting the high-throughput demands of data-intensive operations, particularly when dealing with large-scale datasets and complex computational models. Computational experiments were orchestrated using an internal *SLURM* (Simple Linux Utility for Resource Management) compute cluster. We configured SLURM to efficiently allocate resources based on the demands of queued jobs, with dynamic adjustments based on priority and current load. It should be noted that the computational resources described here sufficed for all phases of the research project; the full project did not require more compute resources than those reported for the experiments.

##### Hyperparameter optimization

We initiated hyperparameter optimization using a randomized grid search approach, applied specifically to the *Afriat* single-cell genomics dataset due to its moderate size, facilitating efficient and thorough exploration of the hyperparameter space. The optimization targeted two primary evaluation metrics: the maximum Mutual Information Gap (maxMIG), serving as a quantitative measure for assessing covariate disentanglement efficiency, and DEG-R^2^, focusing explicitly on reconstruction accuracy for differentially expressed genes. The optimization was conducted through the Weights & Biases (wandb) platform, leveraging *TarDis*’ integrated logging capabilities. Initial random sampling broadly explored hyperparameters across multiple architectural and training-related dimensions, including latent dimensionalities, number of neurons per layer in multilayer perceptrons (MLPs), and learning rate schedules. However, iterative evaluations revealed diminishing returns from broad exploration, leading to focused refinements in a narrower, critical subset of hyperparameters that showed higher sensitivity and impact on model performance. Specifically, hyperparameters such as `weight_decay`, which regulates the extent of L2 regularization thus affecting model complexity, and `kl_warmup_epochs`, influencing the gradual introduction of KL divergence regularization in the Variational Autoencoder (VAE), `use_layer_norm vs use_batch_norm`, and `dropout_rate` were found one of the most impactful ones. Additional sensitivity was observed for hyperparameters defining the strength of individual loss components, which directly modulate the trade-off between reconstruction fidelity and covariate-specific disentanglement. Subsequent to automated grid searches, selective *manual* refinements were performed *occasionally* when configurations surfaced by wandb-tracked experiments demonstrated promising but suboptimal performance, warranting fine-grained adjustments of individual component weights within the auxiliary loss. These adjustments ensured optimal disentanglement without compromising overall model robustness or reconstruction quality.

The auxiliary loss weight *λ*_*C*_ particularly governs the strength of the disentanglement constraints. As seen in Figure 2 and Supplementary Figure 8, increasing *λ*_*C*_ generally improves separation in the reserved latents and boosts clustering metrics like ASW. We did not observe notable overfitting or under-penalization for ranges of *λ*_*C*_ tested on *Afriat*, possibly because that dataset is sufficiently large and the disentanglement constraints are distributed across multiple latent subspaces. We recommend modest grid searches around the default *λ*_*C*_ when working with new datasets of distinctly different sizes or complexity, since an excessively large *λ*_*C*_ could in principle lead to diminished reconstruction if the data are too sparse or limited.

In many single-cell studies, exhaustive hyperparameter tuning is recommended but seldom practiced by the users, complicating reproducibility. By contrast, our aim was to mirror typical real-world usage: researchers often employ a single, default hyperparameter set rather than extensive re-optimizations per dataset. For *TarDis*, once the primary search on *Afriat* was complete, we retained these optimized settings as the default for all subsequent experiments, unless stated otherwise. Notably, this design choice intentionally opts for a more conservative approach; it is likely that further data-specific tuning could yield even better performance on larger or differently structured datasets like *Braun, Suo* and *Norman* datasets.

##### Model and auxiliary loss hyperparameters

**Table 3:** Hyperparameters for model configuration: This table lists the hyperparameters used in the model configuration, including their descriptions and assigned values.

| Parameter | Description | Value |
| --- | --- | --- |
| `n_input` | Number of input features. |  |
| n_batch | Number of batches. If 0, no batch correction is performed. | 0 |
| n_labels | Number of labels. | 0 |
| n_hidden | Number of nodes per hidden layer. Passed into Encoder and Decoder. | 512 |
| n_latent | Dimensionality of the latent space. | $24 + 8 * N_K$ |
| n_layers | Number of hidden layers. Passed into Encoder and Decoder. | 3 |
| n_continuous_cov | Number of continuous covariates. | 0 |
| n_cats_per_cov | A list of integers containing the number of categories for each categorical covariate. | None |
| dropout_rate | Dropout rate. Passed into Encoder but not Decoder. | 0.25 |
| dispersion | Flexibility of the dispersion parameter, which can be "gene", "gene-batch", "gene-label", or "gene-cell", when gene_likelihood is either nb or zinb. | "gene" |
| log_variational | If True, use torch.log1p on input data before encoding for numerical stability (not normalization). | True |
| gene_likelihood | Distribution to use for reconstruction in the generative process. ("zinb", "nb", "poisson") | "nb" |
| latent_distribution | Distribution for the latent space. ("normal", "ln") | "normal" |
| encode_covariates | If True, covariates are concatenated to gene expression prior to passing through the encoder(s). | False |
| deeply_inject_covariates | If True and n_layers > 1, covariates are concatenated to the outputs of hidden layers in the encoder(s) and the decoder. | True |
| batch_representation | Method for encoding batch information. ("one-hot", "embedding") | "one-hot" |
| use_batch_norm | Specifies where to use torch.nn.BatchNorm1d in the model. ("encoder", "decoder", "none", "both") | None |
| use_layer_norm | Specifies where to use torch.nn.LayerNorm in the model. ("encoder", "decoder", "none", "both") | "both" |
| use_size_factor_key | If True, use the anndata.AnnData.obs column as defined by the size_factor_key parameter in the model's setup_anndata method as the scaling factor in the mean of the conditional distribution. | False |
| use_observed_lib_size | If True, use the observed library size for RNA as the scaling factor in the mean of the conditional distribution. | True |
| library_log_means | Vector of shape (1, n_batch) of means of the log library sizes that parameterize the prior on library size. | None |
| library_log_vars | Vector of shape (1, n_batch) of variances of the log library sizes that parameterize the prior on library size. | None |
| var_activation | Callable used to ensure positivity of the variance of the variational distribution. Passed into Encoder. The default is the exponential function. | None |
| deeply_inject_disentangled_latents | If True, deeply inject disentangled latents. | True |
| include_auxillary_loss | If True, include auxiliary loss. | True |
| beta_kl_weight | Weight for the KL divergence term in the loss function. | 0.5 |

**Table 4:**
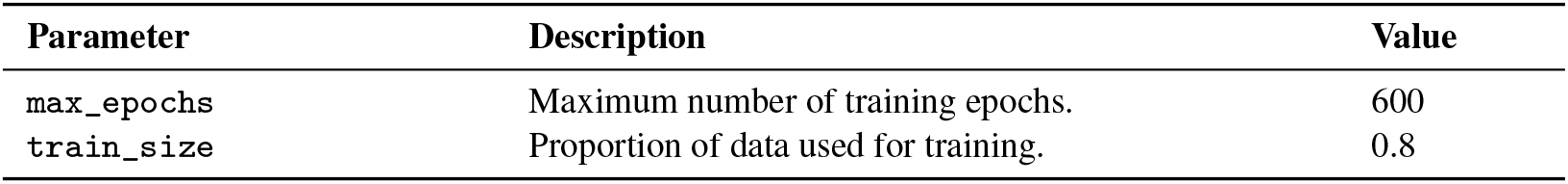
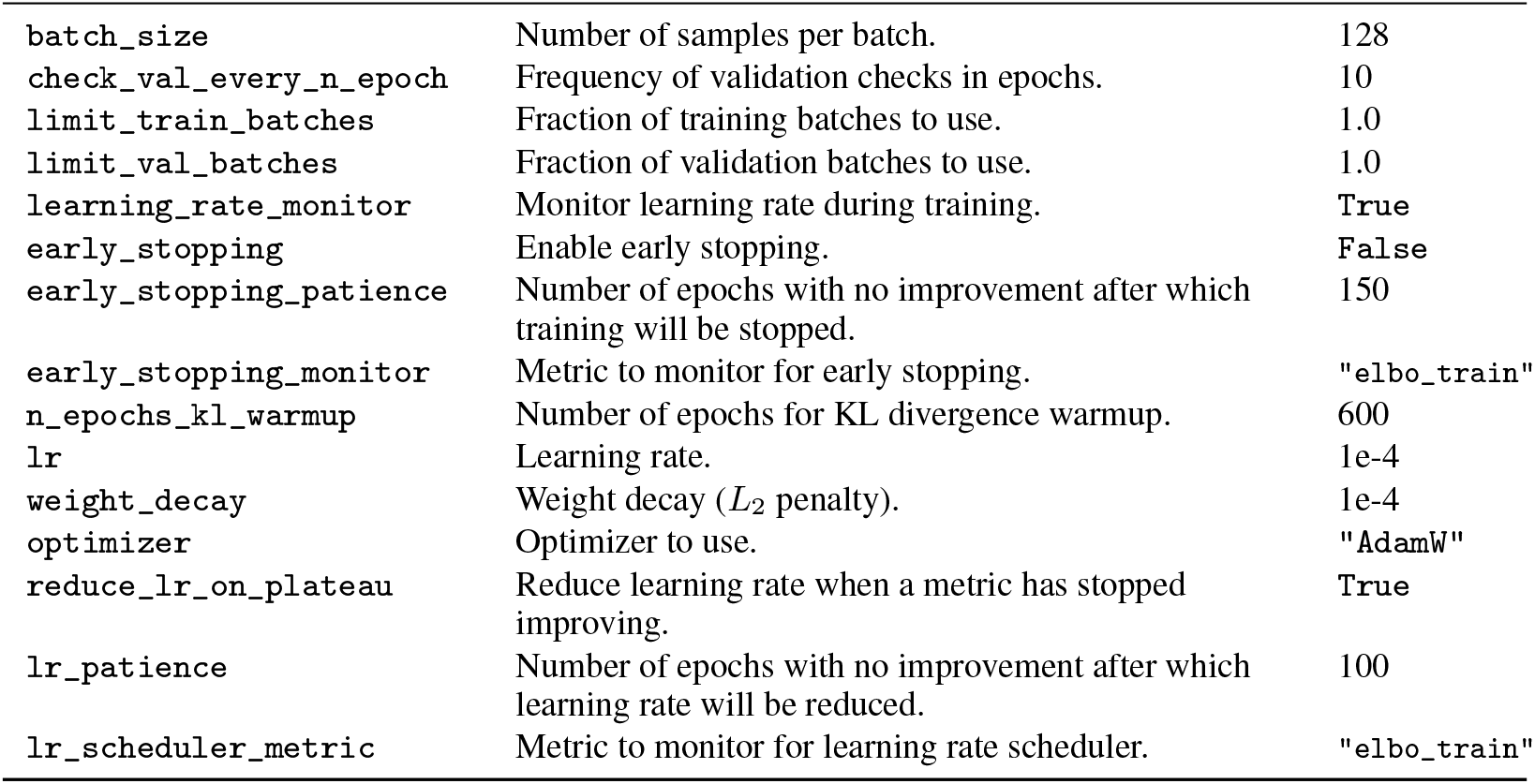
Hyperparameters used for optimization: It provides a comprehensive overview of the configurations necessary to monitor and enhance model performance throughout the training.

| Parameter | Description | Value |
| --- | --- | --- |
| max_epochs | Maximum number of training epochs. | 600 |
| train_size | Proportion of data used for training. | 0.8 |
| batch_size | Number of samples per batch. | 128 |
| check_val_every_n_epoch | Frequency of validation checks in epochs. | 10 |
| limit_train_batches | Fraction of training batches to use. | 1.0 |
| limit_val_batches | Fraction of validation batches to use. | 1.0 |
| learning_rate_monitor | Monitor learning rate during training. | True |
| early_stopping | Enable early stopping. | False |
| early_stopping_patience | Number of epochs with no improvement after which training will be stopped. | 150 |
| early_stopping_monitor | Metric to monitor for early stopping. | "elbo_train" |
| n_epochs_kl_warmup | Number of epochs for KL divergence warmup. | 600 |
| lr | Learning rate. | 1e-4 |
| weight_decay | Weight decay ($L_2$ penalty). | 1e-4 |
| optimizer | Optimizer to use. | "AdamW" |
| reduce_lr_on_plateau | Reduce learning rate when a metric has stopped improving. | True |
| lr_patience | Number of epochs with no improvement after which learning rate will be reduced. | 100 |
| lr_scheduler_metric | Metric to monitor for learning rate scheduler. | "elbo_train" |

**Table 5:**
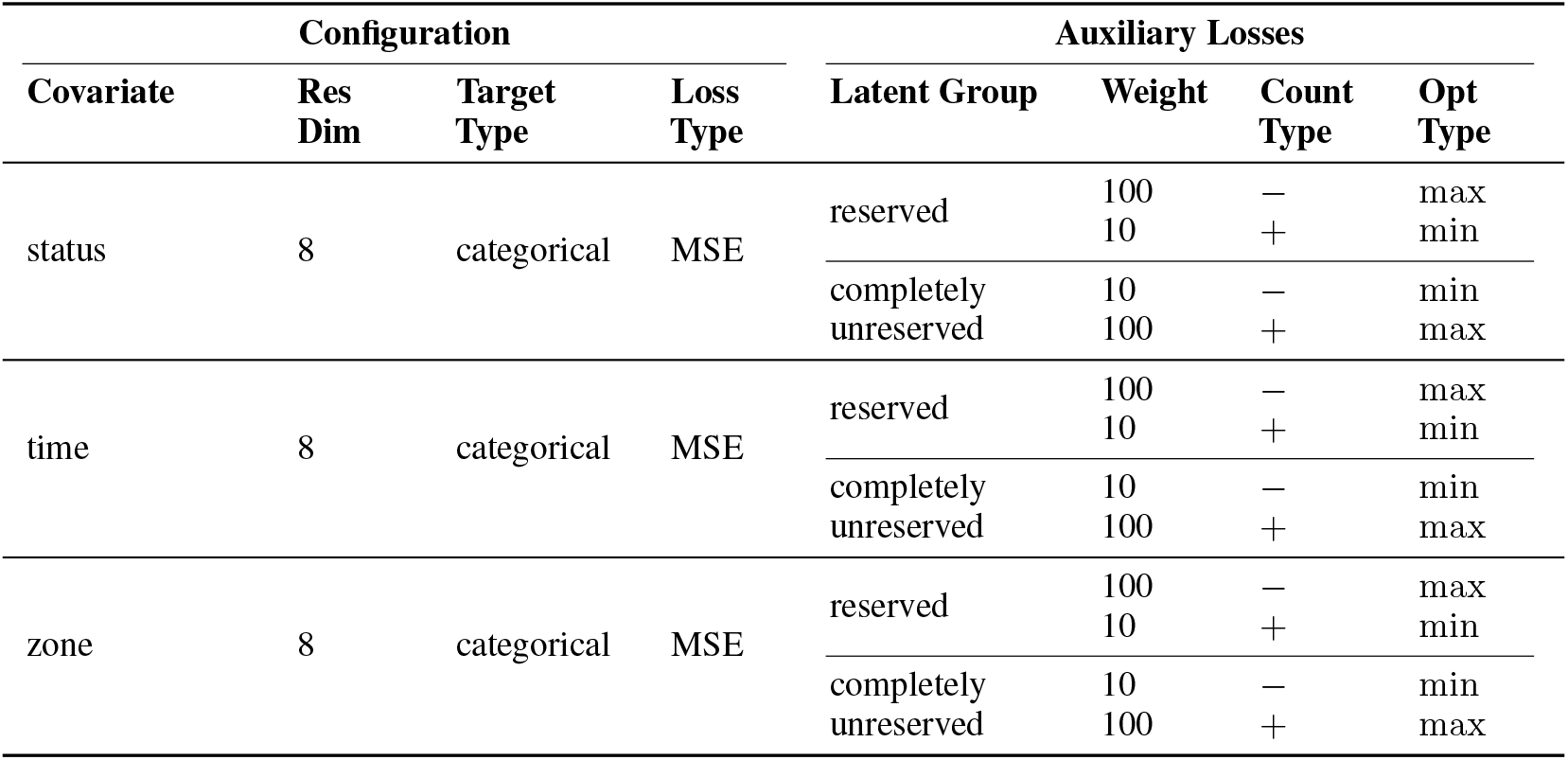
Summary of ℒ_*C*_ configuration designed for covariates, namely *status control, time*, and *zone* in *TarDis*_multiple_ model trained on *Afriat* dataset. It provides insights into how each covariate contributes to the overall model loss.

#### 4.4.4 Theoretical assumptions

- Gene Dependency: The model implicitly assumes that the expression of genes can be considered independently (conditional on the latent space and covariates) when calculating losses. However, genes often exhibit co-expression or are co-regulated, which the model might not account for without specific modifications.
- Homogeneity of Cell Populations: It’s implicitly assumed that cell populations are homogeneous within groups defined by covariates, which might not be the case in heterogeneous biological conditions such as tumors or developing tissues.
- Distribution of Gene Expression Counts: The model assumes that gene expression counts can be modeled effectively using a Negative Binomial distribution. This assumption is common but might not always capture the real variability and distribution in different types of datasets.
- Linearity and Gaussianity of Latent Space: The auxiliary loss assumes a Gaussian distribution for the latent vectors **z**_*nk*_. This implies assumptions about linearity and normality in the latent space, which may not hold in more complex or non-linear biological data structures. This assumption is critical for the model’s simplicity and tractability:

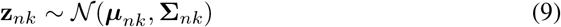
- Static Covariate Definition: The model assumes static and well-defined positive or negative sample definitions in terms of covariate values. This is critical for the stability of the training process: 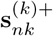 and 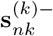 are fixed and consistent throughout the dataset.
- Consistency and Availability of Covariate Labels: Consistent and accurate labeling of covariates across all cells is required. Incomplete or inaccurate labels can undermine the model’s effectiveness:

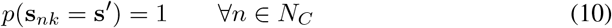
- Smoothness of Latent Space: The auxiliary loss assumes the latent space is smooth and continuous, allowing for meaningful interpolation and extrapolation:

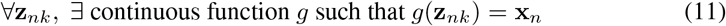
- Sensitivity to Outliers: The model does not explicitly account for outliers, which can skew learned representations. It’s assumed that:

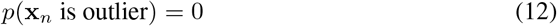
- Assumption of Sufficient Sample Size: The effectiveness of the model in disentangling and accurately representing biological phenomena is contingent upon having a sufficiently large number of samples to cover the variability and complexity of the data. Small sample sizes could lead to overfitting and poor generalization to new data:

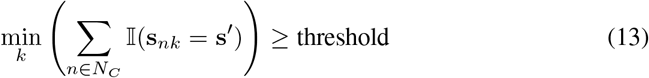
- Data Sparsity: The model assumes it can handle sparsity in single-cell genomic data without additional modifications.
- Consistency of Environmental and Experimental Conditions: It’s assumed that all cells are subject to similar environmental and experimental conditions, aside from the controlled variations represented by covariates. Variability in these conditions could introduce unmodeled noise and bias.

#### 4.4.5 Datasets insights

Please be aware that this section contains embedded hyperlinks, which are essential for accessing the referenced datasets and additional resources. For optimal functionality and ease of navigation, it is highly recommended to consult the PDF version of this document. The PDF format ensures that all hyperlinks are active and can be directly accessed, facilitating seamless retrieval of the associated data and supplementary information.

##### *Afriat* Dataset

The *Afriat* dataset, named after the first author of the study, provides high-resolution single-cell RNA sequencing and single-molecule transcript imaging data of host and parasite gene expression during the liver stage of the rodent malaria parasite *Plasmodium berghei* ANKA. It highlights spatial differences in gene expression across hepatocyte lobule zones, revealing insights into the molecular interactions between host and parasite [1].

*Number of Samples*: The dataset comprises 19,053 individual cells.

*Number of Features*: It encompasses expression profiles across 8,203 genes.

*Source*: The data is publicly accessible. The raw dataset can be found under GEO accession number GSE181725. Processed data are available as a Seurat object [15] at Zenodo. The AnnData [92] format, utilized in this study, was downloaded from Figshare, as prepared by Biolord study [70]. No preprocessing or subsetting was performed on our part.

##### *Suo* Dataset

Named after a co-author of the originating study, the *Suo* dataset offers a multi-organ, single-cell transcriptomic perspective, capturing dynamic immune system developments across nine prenatal human tissues during embryonic stages. This comprehensive dataset details the temporal and spatial maturation of immune cells, highlighting embryonic developmental timing and the interaction between different organ systems in shaping the immune landscape [86].

*Number of Samples*: From an initial count of 908,178 individual cells, 841,922 cells met quality control standards set by established single-cell best practices [33].

*Number of Features*: The dataset, which initially profiled 33,538 genes, has been refined to focus on 8,192 highly variable genes (HVGs), following established single-cell sequencing best practices [33].

*Source*: The raw dataset can be found under ArrayExpress accession number E-MTAB-11343. Processed data are available in AnnData format, accessible at Cellatlas portal. Additional metadata with more detailed annotation is available through the cellxgene server [11]. The metadata was then refined and corrected for errors by the authors.

##### *Braun* Dataset

Named for the first author, the Braun dataset provides a comprehensive single-cell transcriptomic analysis of the human brain during the crucial first trimester. Spanning 5 to 14 postconceptional weeks across 26 brain specimens, the dataset includes over 1.66 million cells dissected into 111 distinct biological samples. This extensive dataset captures the early spatial and transcriptional blueprint of brain development, with detailed insights into neuronal and glial differentiation trajectories [12].

*Number of Samples*: From an initial count of 1,665,937 individual cells, 1,661,498 cells met quality control standards set by established single-cell best practices [33].

*Number of Features*: The dataset, which initially profiled 59,459 genes, has been refined to focus on 8,192 highly variable genes (HVGs), following established single-cell sequencing best practices [33].

*Source*: Raw sequencing data are available from the European Genome Phenome Archive under the accession number EGAS00001004107). The data can be browsed interactively at SciLifeLab Portal and cellxgene server. The metadata was then refined and corrected for errors by the authors.

##### *Miller* Dataset

The Miller dataset, named after the first author of the paper, provides a detailed single-cell mRNA sequencing atlas of human lung development from 11.5 to 21 weeks, integrated with studies on homogeneous human bud tip organoid cultures. This dataset specifically investigates the role of SMAD signaling in the differentiation of bud tip progenitors into airway lineages, showcasing how in vitro conditions mirror in vivo airway structures and function. This comprehensive atlas underscores critical insights into the cellular mechanisms guiding human airway differentiation [63].

*Number of Samples*: From an initial count of 8443 individual cells, 7405 cells met quality control standards set by established single-cell best practices [33].

*Number of Features*: The dataset, which initially profiled 36,601 genes, has been refined to focus on 8,192 highly variable genes (HVGs), following established single-cell sequencing best practices [33].

*Source*: The raw scRNA-seq data associated with this study are available in the EMBL-EBI ArrayEx-press database under accession number E-MTAB-8221. The metadata was then refined and corrected for errors by the authors.

##### *Sciplex* Dataset

The *Sciplex* dataset, derived from the sci-Plex technology using nuclear hashing, quantifies transcriptional responses to chemical perturbations at single-cell resolution. Applied to three cancer cell lines and exposing them to 188 distinct compounds, it evaluates dose-dependent effects and different drug responses. This high-throughput chemical screen profiles approximately 650,000 single-cell transcriptomes across about 5000 samples in a single experiment, revealing cellular heterogeneity in drug response, commonalities within compound families, and nuanced differences within compound types, particularly histone deacetylase inhibitors [84].

*Number of Samples*: The dataset comprises 14,811 individual cells.

*Number of Features*: It encompasses expression profiles across 4999 genes.

*Source*: Both processed and raw data are accessible via NCBI GEO under accession number GSE139944. The dataset used, in its preprocessed and subsetted format, aligns with the methodology described in the CPA paper [56], provided courtesy of the authors of CPA. No further preprocessing or subsetting was conducted by our team.

##### *Norman* Dataset

Named for the first author, the *Norman* dataset leverages high-content Perturb-seq (single-cell RNA-sequencing pooled CRISPR screens) to explore cellular and organismal complexity through combinatorial gene expression. The dataset features transcriptional responses from 284 different single or double gene knockouts, allowing for the exploration of genetic interactions at scale. This includes the mapping of regulatory pathways, classification of genetic interactions such as suppressors, and the mechanistic study of synergistic effects, notably between CBL and CNN1 in erythroid differentiation [66].

*Number of Samples*: The dataset comprises 108,497 individual cells.

*Number of Features*: It encompasses expression profiles across 5000 genes.

*Source*: Raw data is accessible via NCBI GEO under accession number GSE133344. The dataset used, in its preprocessed and subsetted format, aligns with the methodology described in the CPA paper [56], provided courtesy of the authors of CPA. No further preprocessing or subsetting was conducted by our team.

#### 4.4.6 Disentangled Latent Space: Gene Selection Discussion

The approach employed by *TarDis* offers distinct advantages over traditional differential expression (DE) analysis by enabling a deeper, more flexible exploration of gene expression data while addressing technical and biological variability. While DE analysis remains a robust method for identifying gene-covariate associations, it is inherently limited by its reliance on predefined class labels and its susceptibility to information loss when adjusting for confounders. In contrast, *TarDis* leverages covariate-specific latent spaces to uncover nuanced biological patterns, supports causal hypothesis generation, and preserves granularity in continuous covariates. The key advantages include the following:

- Exploratory Analysis within Covariate-Specific Latent Spaces: While differential expression (DE) analysis offers robust insights, *TarDis* enables an exploratory analysis within covariate-specific latent spaces that DE analysis cannot capture. For instance, within a covariate-specific latent, data points can be further clustered into distinct groups, each characterized by unique DE genes. This level of granularity allows for the discovery of subgroups within covariate classes that traditional DE methods, constrained by pre-defined class labels, might overlook. Hence, *TarDis* enables a more nuanced exploration of gene expression dynamics, supporting sophisticated hypotheses generation beyond standard DE analysis. For example, within a certain developmental time point, *TarDis* might identify multiple Leiden clusters within the latent space reserved for *time* for a given cell type. Each of these clusters could represent a subpopulation of cells that, although originating from a similar stage of development, are on divergent paths towards differentiating into distinct cell types. Such subclusters could be crucial for identifying transitional states that are not apparent in traditional DE analysis. This ability allows for the characterization of unique DE genes that define each cluster, providing insights into the molecular mechanisms driving differentiation. In practice, this means that *TarDis* could help researchers uncover previously unrecognized cellular transitions within a homogeneous population, such as identifying progenitor cells in early developmental stages that eventually differentiate into different neuronal subtypes. Each Leiden cluster identified by *TarDis* could potentially highlight a unique pathway of development, characterized by distinct gene expression profiles, thereby offering a more detailed and dynamic view of cellular differentiation.
- Counterfactual Inference: *TarDis* extends beyond identifying associations by enabling counterfactual reasoning within the latent space. By manipulating specific latent dimensions while holding others constant, it is possible to observe the hypothetical outcomes on gene expression, offering a powerful tool for causal inference and understanding the impact of varying covariates in isolation.
- Preservation of Continuous Covariates Beyond Discretization: Traditional DE approaches often require the discretization of continuous covariates (e.g., age, dosage), potentially leading to loss of granularity and introducing biases. In contrast, *TarDis* maintains the integrity of continuous variables through its latent representations, employing a distance-weighted loss function that accurately captures subtle biological shifts, thus providing a richer and more precise characterization of gene expression changes over continuous covariates.
- Minimizing Artifact Propagation: Standard DE analyses can inadvertently adjust for batch effects in a manner that either strips away genuine biological signals or over-corrects, leading to skewed gene expression profiles. *TarDis* addresses this by explicitly modeling technical artifacts within distinct latent dimensions, thereby preserving true biological variability in the primary latent space reserved for invariant features. This targeted segregation helps in maintaining the purity of biological insights derived from the data.
- Scalability and Applicability to Multi-Omics Data: Considering the increasing size of single-cell datasets, *TarDis*’s architecture is designed to scale linearly with the number of cells and genes, facilitating efficient processing of large-scale data. Additionally, the flexibility of the underlying variational autoencoder framework makes *TarDis* adaptable to various omics data types, providing a unified approach for complex analyses that would be cumbersome with traditional DE strategies.

### 4.5 Quantification and statistical analysis

#### 4.5.1 Loss functions

Without loss of generality, various choices for the loss function are investigated, focusing on elucidating the loss incurred between the anchor point **x**_*nk*_ and the positive sample 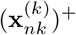. The loss between the anchor point and the negative sample 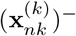 can be derived similarly, with appropriate adjustments to maximize this loss.

##### Mean Squared Error (MSE)

The MSE between the latent representation of the anchor **z**_*nk*_ and its positive counterpart 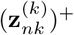 for the *k*th covariate is given by:

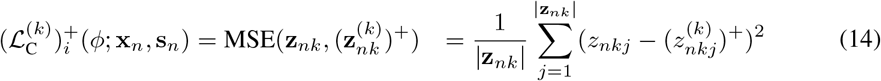

However, minimizing the *L*_2_ distance between normal vectors from distinct multivariate normal distributions with unique diagonal covariance matrices does not inherently ensure the convergence of their distributions. While this minimization may align distribution means, it disregards differences in variances and higher-order moments essential for comprehensive distributional characterization.

Mathematically speaking, if **z**_*nk*_ ~ 𝒩(***µ***_*nk*_, **Σ**_*nk*_) and 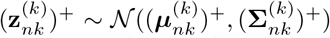, by using linearity of expectation and properties of the transpose, the expected squared *L*_2_ distance between **z**_*nk*_ and 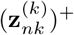 can be simplified to:

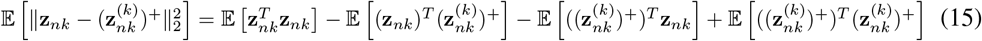

For any vector **z**_*nk*_ with mean ***µ***_*nk*_ and covariance **Σ**_*nk*_, the following identity holds:

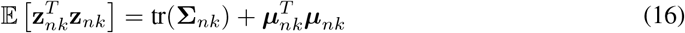

Applying this to 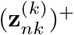 and also knowing **z**_*nk*_ and 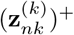 are independent, we have:

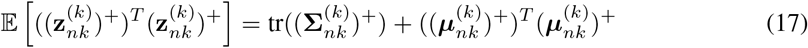

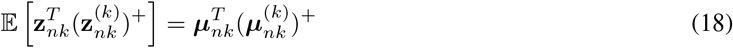

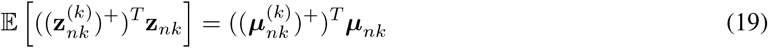

where tr(·) denotes the trace of a matrix. Substituting back, we find:

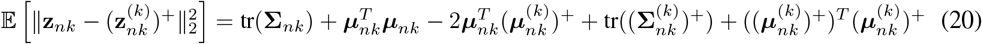

To simplify further, recognizing the vector identity 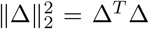 for squared terms where 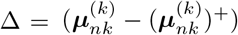:

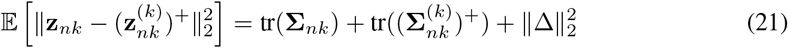

This expression reveals that the expected squared *L*_2_ distance depends on both the aggregate covariances and the squared difference between the means. Minimizing this distance reduces the mean disparity term 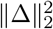, but does not necessarily minimize the covariance term tr 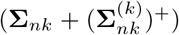, which reflects distributional variability. However, it is crucial to ensure the convergence of our latent representations of similar pairs across their entire characteristics. Notably, as Tong and Kobayashi [87] demonstrated, differences in the diagonal covariances of multivariate normal distributions can significantly influence the optimal transport cost and Wasserstein distance, even when the means are aligned. This highlights the importance of considering both mean and covariance differences for accurate distribution comparison. Consequently, we redirect our focus towards statistical metrics like KL divergence, which encompass the entire distribution and provide a more comprehensive assessment of distributional convergence.

##### KL Divergence

Unlike the *L*_2_ distance, which primarily measures central tendency, the KL divergence accounts for both dispersion and correlation structure. Specifically, KL divergence is sensitive to differences in the means and covariance matrices of the distributions, offering a comprehensive measure of how well one distribution approximates another, beyond merely the distance between their centers.

To frame our problem contextually, assume we have determined the representation of a positive data point in a lower-dimensional space, i.e., 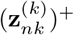 is fixed. With this in mind, we aim to represent the anchor point to reflect its partial similarity in its corresponding latent representation **z**_*nk*_. Therefore, we utilize the encoder distribution of the positive sample, 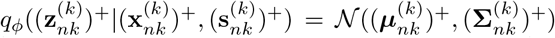 as the target for the current point’s distribution, *q*_*ϕ*_(**z**_*nk*_ | **x**_*nk*_, **s**_*nk*_) = 𝒩 (***µ***_*nk*_, **Σ**_*nk*_) given that the gradients for the forward pass of the positive sample are not computed.

Based on the KL divergence between these two multivariate Gaussian distributions, the positive pair loss 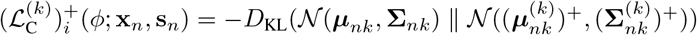 can be calculated using a straightforward and efficient formula:

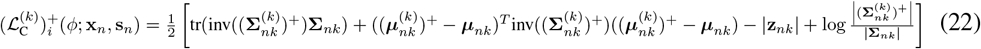

Here, inv(.) stands for the inverse of a matrix, |.| represents the determinant of a matrix, |**z**_*nk*_| is the dimensionality of the distributions, 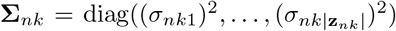 and 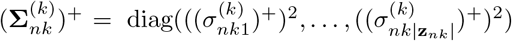. Furthermore, the determination of the determinant for such matrices is simplified, requiring only the multiplication of their diagonal elements. Therefore, equation 22 becomes:

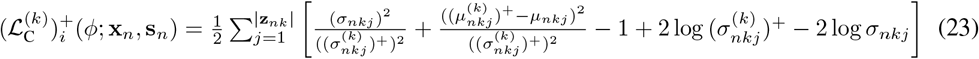

We propose summing the KL divergence over all covariates *k*, analogous to the total correlation (TC) in the objective function of the Relevance Factor VAE (RF-VAE)[44]. This approach is designed to promote independence among latent variables. Consequently, we apply this method to the KL loss term by calculating the KL divergence between each latent representation and the standard normal distribution individually, and then summing the results.

Additionally, instead of assigning a weight to each positive pair loss function with respect to covariate *k* and the KL divergence between its latent representation and the prior distribution (standard normal distribution), we introduce relevance indicators, **r**^(*k*)^ and 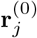 respectively. These indicators can be learned via a variational approach. They are parameterized and updated during the training process.

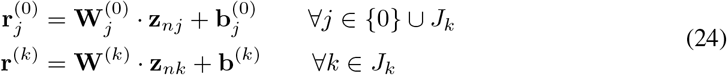

Hence the primary objective function to maximize for becomes:

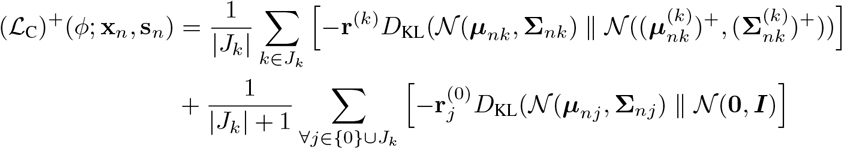

##### Bhattacharyya Loss

When comparing the Bhattacharyya Loss (*D*_*B*_) to the KL divergence, several key distinctions arise. KL divergence can be less effective in handling outliers and noise compared to *D*_*B*_, which provides a more robust measure in noisy environments [81]. Studies have demonstrated that in high-dimensional data scenarios, *D*_*B*_ can outperform KL divergence in both clustering accuracy and robustness to data anomalies [17].

Incorporating *D*_*B*_ as a loss function offers several additional advantages. First, it has shown superior performance in distinguishing between different distributions, which is essential for effective novelty detection [82] and a key aspect of disentanglement. Disentangling different factors of variation in the data often requires a measure that can accurately differentiate between various underlying distributions. Thus, the superior performance of *D*_*B*_ in this regard directly supports its use in disentanglement tasks. In the domain of single-cell RNA sequencing (scRNA-seq), *D*_*B*_ has been successfully applied to detect fear-memory-related genes from neuronal data, demonstrating its ability to handle the high heterogeneity and dropout noise inherent in such datasets [101]. Furthermore, it has been integrated into k-means clustering, enhancing the efficiency and memory-saving capabilities for large-scale scRNA-seq data analysis [8]. *D*_*B*_ is also robust to outliers and noise, ensuring more reliable and consistent results, which is crucial for noisy datasets [64]. Disentangling factors of variation in noisy datasets requires a measure that can reliably handle outliers and noisy data points without compromising the integrity of the disentangled components. *D*_*B*_’s robustness makes it a suitable choice for such tasks. Additionally, its symmetry and comprehensive capture of distributional differences enhance the accuracy of various analytical models [94]. For disentanglement, accurately capturing and separating the underlying factors of variation in the data is essential. *D*_*B*_’s mathematical properties ensure that it can provide a more precise and reliable measure of these differences, facilitating better disentanglement.

Therefore, we can write the positive pair loss utilizing *D*_*B*_ as follows:

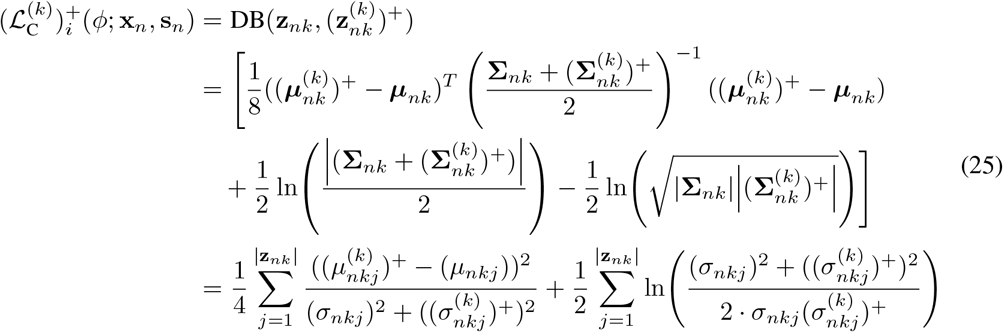

##### Mahalanobis Loss

Mahalanobis Loss (*D*_*M*_) is a robust metric for quantifying the distance-like measure between a point and a distribution, or between two points within a distribution-defined space. Unlike KL divergence and *D*_*B*_, *D*_*M*_ measures the deviation of a point from the mean of a distribution and can be extended to compare the central tendencies of two distributions.

The innovative use of *D*_*M*_ significantly enhances data interpretation and clustering accuracy. The DR-A model, combining a VAE with a generative adversarial network (GAN) leverages *D*_*M*_ for dimensionality reduction, achieving superior clustering and more precise low-dimensional representations of scRNA-seq data [53]. This precision is crucial for accurately representing covariates in lower-dimensional spaces.

The scDREAMER framework integrates *D*_*M*_ within an adversarial VAE to tackle skewed cell types and nested batch effects, improving batch correction and preserving biological variability across heterogeneous datasets [79]. Table 1 highlights that while our model excels in batch correction, there is room for improvement in biological conservation. Therefore, we can adopt *D*_*M*_ to measure the dissimilarity between the latent representation of the anchor point **z**_*nk*_ and the respective posterior distributions 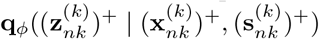as follows:

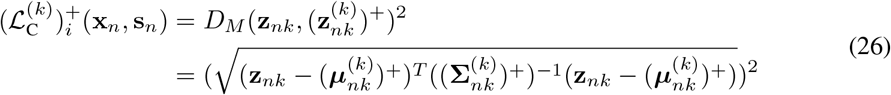

The inverse covariance matrix computation simplifies to the reciprocal of each diagonal element, resulting in:

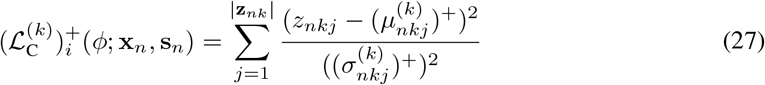

Minimizing *D*_*M*_ encourages **z**_*n*_ and 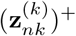 to be located within high-probability regions of the latent space, as defined by the Gaussian distribution. The latent representation of the positive example 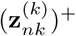 serves as a reference, with all adjustments made relative to the current anchor point **z**_*nk*_.

##### Fisher Information

Fisher information can be used to measure the amount of information that a random variable 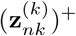carries about the unknown parameters ***µ***_*nk*_ and **Σ**_*nk*_ of a probability distribution modeling 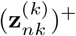. This measurement allows for a more precise identification of the most informative latent factors, leading to more interpretable representations. Because Fisher information is grounded in information theory, the resulting disentangled factors are often more meaningful and easier to understand, which is beneficial for tasks requiring human interpretability of covariates [89]. Representations derived using Fisher information have been shown to improve performance in downstream tasks such as classification, clustering, and anomaly detection [42], which is the ultimate goal of learning latent representations of single-cell RNA-seq data. Therefore, in the context of VAEs, Fisher information aids in analyzing information loss during the encoding process:

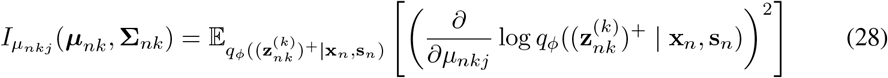

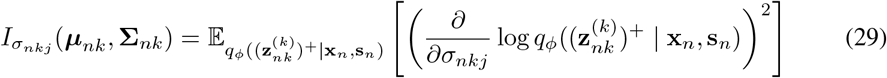

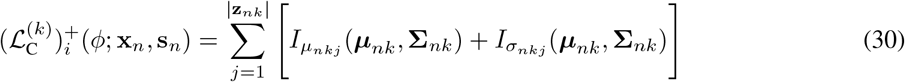

In our case, the log-likelihood function for a single observation **x**_*n*_ is given by:

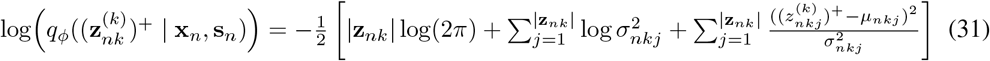

For the mean parameter *µ*_*nkj*_:

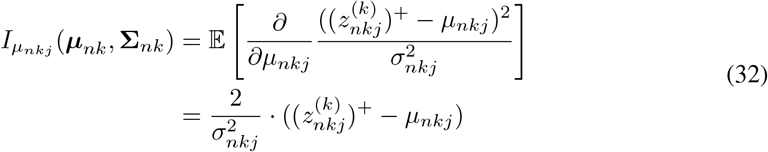

For the variance parameter 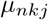:

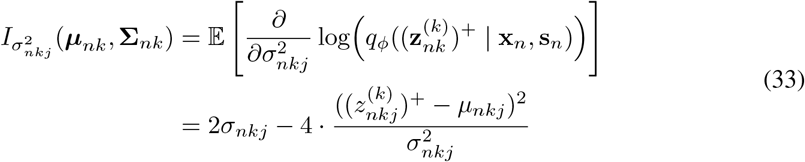

#### 4.5.2 Evaluation metrics

##### Average Silhouette Width

The average silhouette width (ASW) [75] evaluates clustering quality by measuring the relationship between within-cluster and between-cluster distances. ASW values range from −1 to 1, where −1 indicates misclassification, 0 indicates overlapping clusters, and 1 indicates well-separated clusters.

For each data point **x**_*n*_, the silhouette coefficient s(**x**_*n*_) is calculated as:

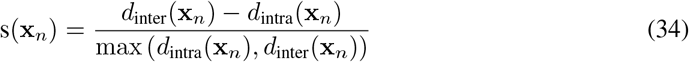

where *d*_intra_(**x**_*n*_) is the average distance from point **x**_*n*_ to all other points within the same cluster (intra-cluster distance) and *d*_inter_(**x**_*n*_) is the minimum average distance from point **x**_*n*_ to points in any other cluster (nearest-cluster distance). The overall ASW is the mean of the silhouette coefficients for all points in the dataset:

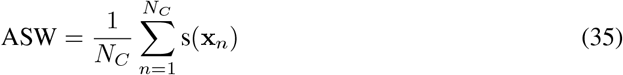

where *N*_*C*_ is the total number of data points. ASW is particularly relevant in single-cell genomics for assessing how well cells cluster based on their gene expression profiles [74]. This metric provides an intuitive measure of clustering quality and batch mixing, crucial for understanding both biological conservation and batch effect removal. It is particularly useful in clustering-based analyses but may be sensitive to noise and outliers.

##### Cell Type Average Silhouette Width

Cell type average silhouette width (Cell type ASW) [60] evaluates cell clustering quality in single-cell transcriptomics by measuring how well cells are grouped based on type labels. The silhouette coefficient for each cell is computed similarly to general ASW. To scale the ASW values between 0 and 1, the following transformation is applied:

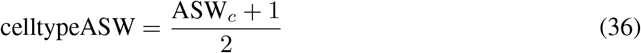

where ASW_*c*_ is the ASW computed over all cell type labels *c*.

##### Batch Average Silhouette Width

Batch average silhouette width (Batch ASW) [60] assesses the quality of batch mixing in integrated datasets, which is essential in single-cell transcriptomics to ensure that technical variations do not obscure biological signals. The silhouette coefficient for each cell, based on batch labels, is computed similarly to general ASW.

To obtain a Batch ASW score between 0 and 1, the following transformation is applied for each batch label *j*:

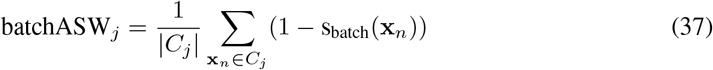

where *C*_*j*_ is the set of cells with batch label *j*, |*C*_*j*_| is the size of this set, and s_batch_(*n*) is the silhouette coefficient for each cell *n* based on batch labels. The final Batch ASW score is calculated by averaging the batch ASW values across all batch labels:

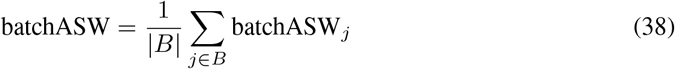

where *B* is the set of unique batch labels. A Batch ASW score closer to 0 indicates good batch mixing, meaning batch effects have been effectively corrected [29].

##### Isolated Label F1 Score

Precision, also known as positive predictive value, gauges the proportion of correctly predicted positive instances among the total predicted positives. It’s calculated by considering True Positives (TP) against False Positives (FP), following the formula:

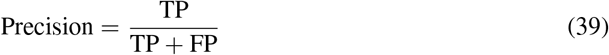

In contrast, Recall, also called sensitivity or true positive rate, measures how well the model identifies actual positive instances, crucial when false negatives are costly. Its calculation focuses on TP relative to FN, given by:

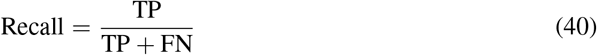

The F1 score, a harmonic mean of precision and recall, offers a single metric balancing both aspects, with high values indicating a well-balanced model. It is calculated as:

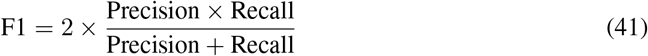

Isolated Label Scores are used to evaluate the clustering and separation of cell identity labels shared by a few batches. Specifically, the isolated label F1 score, also known as the class-wise F1 score, evaluates the F1 score for individual classes and is optimized to achieve the best clustering of these isolated labels, ensuring effective integration of rare cell types. This metric is particularly valuable for handling imbalanced datasets, such as those in single-cell genomics, where it assesses the accuracy of identifying rare cell types [60, 83]. The original scIB package typically employs a cluster-based F1 scoring method by default. However, for the sake of speed and simplicity, we are opting to use the ASW instead as implemented in `scib-metrics` package [26]. The isolated label ASW measures the separation quality of these labels. These scores address the challenge of integrating rare cell types, ensuring that integration methods can effectively manage rare cell populations. However, the performance of these scores is heavily influenced by the quality of initial annotations.

##### Mutual Information

Mutual information (MI) quantifies the reduction in uncertainty about one variable given knowledge of another between variables in complex systems, making it a valuable measure in both theoretical analyses and practical applications [23, 50]. It measures the amount of information shared between two random variables 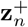 and 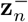 as follows:

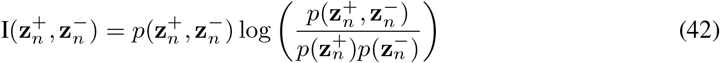

where 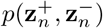 is the joint probability distribution of 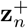 and 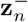 and 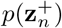 and 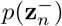 are their marginal distributions.

The value of MI is non-negative, 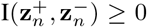, and measures the reduction in uncertainty of 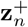 given 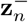 and vice versa. When 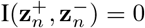, the variables are statistically independent, meaning that knowing 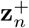 does not provide any information about 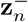. A higher value of MI indicates a greater level of dependency between the variables.

##### Normalized Mutual Information

MI is influenced by dataset size and cluster entropy, complicating comparisons across datasets. Normalization techniques, which adjust MI to a standard range, typically [0, 1], enable more equitable comparisons.

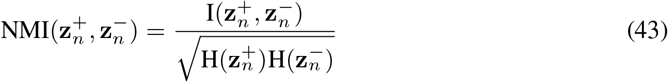

where 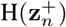 and 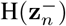 are the entropies of 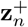 and 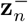.The higher values indicate superior clustering quality [91]. In the context of single-cell genomics, the normalized mutual information (NMI) is critical for evaluating how well clusters correspond to known cell types [60]. This metric evaluates how well cell-type labels are preserved post-integration. It is often used in scenarios requiring validation of clustering results against known labels. While it provides an intuitive measure, it may not distinguish well between near-perfect and perfect clustering.

In response to concerns about the relatively low ARI values (below 0.3) in Table 1, especially when contrasted with the higher NMI scores (above 0.6), which may raise questions about the models’ overall performance, we offer the following clarifications:

- Biological Relevance Despite Lower ARI: While ARI values are influenced by cluster granularity, the key biological groups (e.g., cell types or states) remain discernible in the integrated data. For example, in our case studies (Section 3.2–3.3), marker gene analysis and UMAP visualizations confirm that major cell populations are well-separated. The lower ARI reflects technical discrepancies in cluster counts rather than a failure to capture biological signals. Thus, the integration quality is sufficient for downstream tasks like differential expression or trajectory inference, which rely on meaningful biological variation rather than strict alignment with predefined annotations.
- Default scIB Clustering Settings: In our workflow, we adhere to the default scIB parameters (of `scib_metrics` implementation [25]) and do not manually optimize the number of clusters for each approach. Our focus is to provide a fair, consensus-oriented comparison of different data integration methods, rather than fine-tuning each model to achieve the best possible clustering outcome. Tuning these parameters individually could raise concerns of overfitting and is also *not* the primary aim of this study, which is to investigate *TarDis*’s efficacy in disentangling covariates and preserving biological signals. Importantly, in scIB, the interest lies predominantly in the relative ranking across different models rather than absolute metric values. Consequently, our key takeaway is the comparative improvement in capturing meaningful cell-group distinctions rather than the absolute level of cluster-label alignment.
- Clustering Metrics and Biological Interpretability: Although ARI values near 0.3 may appear lower in comparison to NMI above 0.6, we find that such an ARI range is frequently observed when the number of predicted clusters differs significantly from the number of annotated categories in single-cell data. ARI heavily penalizes discrepancies in the cardinality of clusters, potentially magnifying even small misalignments between predicted clusters and ground-truth labels. We note that default settings in the scIB metrics pipeline often lead to higher cluster counts, which can naturally deflate ARI values.
- Biological Validation Beyond ARI: While ARI is a recognized external clustering metric, our analyses also involve UMAP visualizations and additional single-cell integration assessments (such as silhouette width, LISI score, and more). These complementary metrics more holistically capture how effectively a method corrects batch effects while retaining crucial biological variation.

##### Maximum Mutual Information Gap

The maximum mutual information gap (maxMIG) is a metric designed to evaluate the disentanglement of latent variables in complex datasets where the number of covariates exceeds two, a complexity that only particular methods are equipped to manage [19, 34, 43, 51, 96] due to its ability to generalize and be unbiased [19, 56, 76]. This measure quantifies the MI between latent representations and observed covariates, focusing on how effectively these latent variables independently capture the informative characteristics of each covariate.

The maxMIG is defined for a set of latent variables 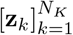 and corresponding covariates 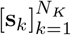 as:

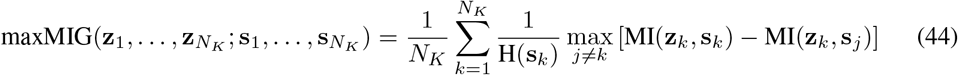

The maxMIG score is computed by averaging the normalized differences between the mutual information of each latent variable with its corresponding covariate and the highest mutual information with any other covariate. This focus on maximizing the information gap helps evaluate the specificity and relevance of each latent variable to its respective covariate. Higher maxMIG values suggest better disentanglement, indicating that each latent variable is more uniquely aligned with a specific covariate, thus enhancing the model’s interpretability and generalizability.

##### Rand Index

The Rand index (*RI*) serves as a pivotal metric for evaluating the concordance between two clustering outcomes. It quantifies the degree of similarity by scrutinizing the allocation of data points into clusters across two distinct clustering results. Computed as the ratio of the sum of agreements to the total number of data point pairs, *RI* encapsulates both intra-cluster cohesion and inter-cluster separation. The formula for calculating the Rand Index is as follows:

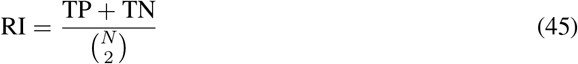

where *N* = TP + TN + FP + FN. While the Rand Index offers valuable insights into clustering performance, it may have limitations when dealing with varying cluster sizes or datasets with an uncertain number of clusters.

##### Adjusted Rand Index

The RI quantifies the proportion of agreements between the two clusterings out of all possible pairings of elements. However, because the RI does not adjust for the chance grouping of elements, the Adjusted Rand Index (ARI) [38, 60] is often preferred, which is defined as:

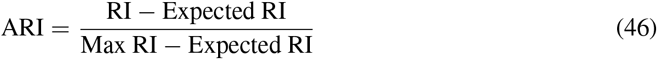

where the Expected RI is the expected value of the RI for random clusterings and the Max RI is the maximum possible value of the RI. Mathematically, the ARI can be expressed as:

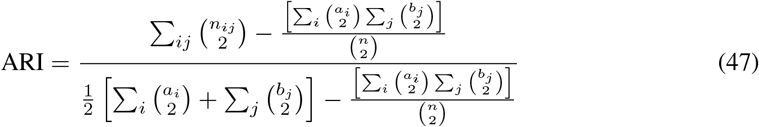

where *n*_*ij*_ is the number of elements in the intersection of cluster *i* in *X* and cluster *j* in *Y, a*_*i*_ is the number of elements in cluster *i* of *X, b*_*j*_ is the number of elements in cluster *j* of *Y*, and 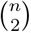 denotes the binomial coefficient. This adjustment provides a corrected-for-chance measure, making the ARI a more reliable metric for clustering comparison.

Values of ARI above zero indicate better-than-random agreement, with a value of 1 representing perfect agreement [39]. In single-cell data analysis, ARI is useful for validating the consistency of cell type assignments across different clustering methods. This metric is key for evaluating clustering performance in the presence of noise and is commonly used to validate clustering results in datasets with known ground truth. However, it can be less intuitive to interpret compared to simpler metrics.

##### k-nearest neighbor Batch Effect Test

The k-nearest neighbor batch effect test (kBET) [16, 60] assesses batch effects in high-dimensional datasets by testing the homogeneity of batch labels within the k-nearest neighbors of each data point. It evaluates whether the neighbors of a cell are more likely to come from the same batch than expected under random mixing. kBET is a robust method designed to quantify batch effects in single-cell RNA sequencing (scRNA-seq) data. To implement kBET, one first constructs a k-nearest-neighbor (kNN) graph for each cell in the dataset, using an appropriate distance metric such as Euclidean distance in a principal component analysis (PCA)-reduced space. For each cell *n*, the algorithm identifies its *k* nearest neighbors and calculates the proportion of cells from each batch within this neighborhood, denoted as 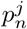, where *j* indexes the batches. Under the null hypothesis of no batch effect, the expected proportion of cells from each batch should reflect the overall batch composition in the dataset, represented as *q*_*j*_. The kBET then compares the observed batch proportions 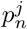 with the expected proportions *q*_*j*_ using a statistical test, such as the Chi-square test or a permutation-based test. The test statistic for each cell *n* is computed as

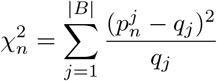

where |*B*| is the number of batches. The p-value associated with the Chi-square statistic indicates the likelihood that the observed batch composition within the neighborhood of cell *n* is consistent with the global batch composition. These p-values are aggregated across all cells to assess the overall presence of batch effects in the dataset. The kBET statistic is:

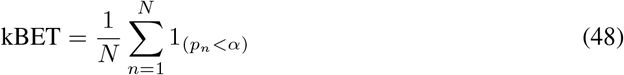

where *N* is the number of neighborhoods tested, *p*_*n*_ is the p-value from a chi-squared test, and *α* is the significance threshold.

This method was evaluated using peripheral blood mononuclear cells (PBMCs) from healthy donors, effectively distinguishing cell-type-specific inter-individual variability from changes in relative proportions of cell populations. kBET is crucial for evaluating the effectiveness of batch effect correction methods in single-cell transcriptomics. The kBET tool and its detailed implementation are available on the kBET GitHub repository.

##### Graph Connectivity

Graph connectivity evaluates whether the kNN graph of integrated data effectively connects all cells with the same identity. For each cell identity label, a subset kNN graph is created. The graph connectivity score is then computed as the average size of the largest connected component relative to the number of nodes with that cell identity [60]. This metric ensures that cells of the same type remain connected post-integration, a critical aspect for evaluating graph-based methods. Despite its importance, calculating graph connectivity can be computationally intensive for large datasets.

In single-cell genomics, graph connectivity assesses the robustness of cell interaction networks. The formula for graph connectivity is:

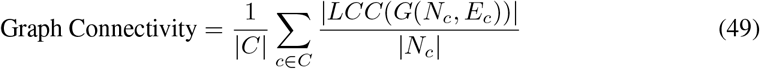

where *C* is the set of cell identity labels, LCC(*G*(*N*_*c*_, *E*_*c*_)) is the largest connected component of the graph for cells with label*c*, and|*N*_*c*_| is the number of nodes with cell identity *c*.

##### Coefficient of determination in VAE

The *R*^2^ Reconstruction metric, often referred to as the coefficient of determination, is a statistical measure used to evaluate the performance of VAEs in reconstructing input data. This metric quantifies how well the reconstructed outputs from a VAE approximate the original inputs, indicating the proportion of variance in the data that is captured by the model. *R*^2^ Reconstruction is particularly useful in the evaluation of VAEs because it provides a clear metric to gauge the accuracy of data reconstructions, facilitates comparison between different VAE architectures or configurations on the same dataset, helps identify areas where the model might be lacking, guiding further refinements. This metric is critical for researchers and practitioners using VAEs to ensure that their models not only generate new data that is statistically similar to the input data but also effectively reconstruct specific instances of input data [32, 41].

In the context of VAEs, the *R*^2^ Reconstruction is defined as:

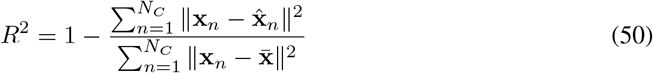

where **x**_*n*_ represents the original input data, 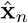 represents the reconstructed data produced by the VAE, and 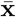 is the mean of the original input data.

The *R*^2^ value ranges from 0 to 1, where a higher value indicates that the model has effectively captured more of the variance in the input data through its reconstructions. An *R*^2^ value of 1 signifies perfect reconstruction, whereas a value close to 0 indicates that the model performs no better than a model that would simply predict the mean of the input data for all outputs.

##### Coefficient of determination for Differentially Expressed Genes in VAE

In computational biology, the evaluation of VAEs reconstruction often focuses on differentially expressed genes (DEG), which show significant changes in expression under different conditions, are critical for understanding biological processes and disease mechanisms. The *R*^2^ Reconstruction metric is adapted in this context to specifically assess how well VAEs can reconstruct the expression patterns of these DEG. Refer to Section 4.5.2 for details of *R*^2^ reconstruction score [32, 41].

The *R*^2^ Reconstruction for DEG is defined as:

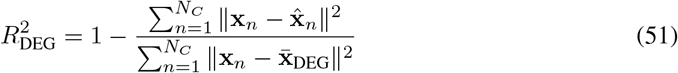

where **x**_*n*_ represents the expression levels of DEG in the original data, 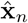 represents their reconstructed levels from the VAE, and 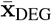 is the mean expression level of DEG.

Focusing on DEG, the *R*^2^ Reconstruction metric specifically evaluates how effectively the VAE captures the variability and regulatory patterns in gene expression that are most biologically relevant and likely to be impacted by experimental conditions. A high *R*^2^ value indicates that the VAE has effectively learned to model the critical aspects of gene expression relevant to the study’s goals.

Reconstructing differentially expressed genes is inherently more difficult yet more critical than reconstructing overall gene expression due to several factors:

(i) *Biological Relevance* DEG often carry more biological significance than stably expressed genes, directly reflecting the cellular responses to biological stimuli or disease states.
(ii) *High Variability* DEG typically exhibit high variability in expression levels, making accurate reconstruction a complex challenge that tests the model’s sensitivity and precision.
(iii) *Data Reduction* By concentrating on DEG, researchers can reduce the dimensionality of the data, focusing computational resources and analytical efforts on the most informative parts of the dataset.
(iv) *Improved Sensitivity* Models tuned to capture changes in DEG can be more sensitive to subtle but biologically important changes that might be overlooked when considering all genes.

Evaluating VAE performance using the *R*^2^ Reconstruction metric on DEG provides insights into the model’s ability to handle the most critical and dynamic components of biological data, facilitating the development of more accurate and biologically informative models.

##### Principal Component Regression

The principal component regression (PCR) quantifies batch removal by calculating the variance contribution of the batch effect per principal component (PC) [60]. The variance contribution of the batch effect is computed as the product of the variance explained by each PC and the corresponding *R*^2^ value from a linear regression of the batch variable onto each PC. Mathematically, it is expressed as:

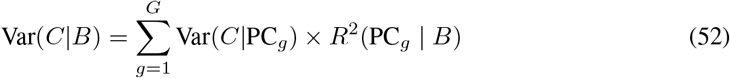

where Var(*C*|PC_*g*_) is the variance of the data matrix *C* explained by the *g*th principal component and *R*^2^(PC_*g*_|*B*) is the coefficient of determination for the batch variable *B*. This metric provides a quantitative measure of batch effects, allowing for direct comparison between methods, and is essential for assessing how well integration methods remove technical variability, particularly in large-scale multi-batch studies. However, it may not fully capture non-linear batch effects.

##### Local Inverse Simpson’s Index

The graph local inverse Simpson’s index (LISI) is a metric for evaluating batch mixing (iLISI) and cell-type separation (cLISI) in integrated single-cell datasets. It uses graph-based distances and the inverse Simpson’s index to measure diversity within neighborhood compositions. Scores are rescaled from 1 to the total number of batches to a range of 0 to 1, where 0 indicates minimal integration or separation, and 1 indicates optimal mixing or segregation. This metric is especially useful for graph-based integration methods and allows for cross-method comparisons, although it requires careful parameter tuning and interpretation [49, 60].

cLISI assesses the integration of diverse cell types within a combined dataset. For each cell, its kNN are identified, and the composition of cell types within this neighborhood is analyzed. The diversity is quantified using the Inverse Simpson’s Index:

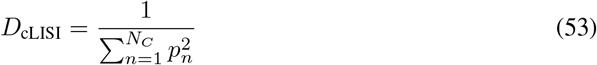

where *p*_*n*_ is the proportion of the *n*-th cell type in the neighborhood, and *N*_*C*_ is the total number of distinct cell types. The average cLISI score across all cells indicates how well cell types are mixed, with high values showing effective mixing and low values indicating poor mixing.

iLISI measures dataset mixing within the local neighborhood of each cell, quantifying how well cells from different datasets are integrated. iLISI close to the number of datasets suggests good mixing, meaning datasets are well integrated where cLISI close to 1 indicates good preservation of cell types, meaning different cell types remain well separated.

Balancing iLISI and cLISI ensures datasets are integrated effectively while preserving distinct cell type identities. Graph LISI’s unified measure for both batch mixing and cell-type separation makes it a valuable tool for single-cell data integration studies, providing a standardized framework for comparing integration methods and identifying optimal strategies.

##### CI calculation and DE gene selection procedure

Steps and rationale for preparing Figure 3.

- Gene Expression Smoothing: To reduce noise and capture underlying trends, each gene’s expression profile along the continuous latent variable (PC1) was convolved with a Gaussian kernel. The kernel’s window size (256 samples) and standard deviation (256) were chosen to balance resolution and smoothness, ensuring local variations are preserved while suppressing high-frequency noise. This generates a moving average trajectory for each gene.
- Confidence Interval Construction: A symmetric confidence interval (CI) around zero was derived under the null hypothesis of no differential expression. Assuming Gaussian-distributed noise in the smoothed expression values, the CI width was determined by computing the cumulative distribution function (CDF) at *±*2.25 standard deviations. This corresponds to a ~ 97.5% two-tailed probability coverage (i.e., CI = 2 *×* Φ(2.25) *−* 1*≈* 0.975), where Φ is the standard normal CDF. Genes with trajectories remaining within *±*1.9 across the latent axis were classified as non-DE (within CI).
- Threshold-Based DE Detection: Genes were deemed differentially expressed (DE) if their smoothed trajectory exceeded the *±*1.9 threshold at any point along the latent axis. This threshold corresponds to the 97.5% quantile under the null distribution, controlling the false positive rate (FPR) at ~ 2.5% per tail. The threshold was applied post-smoothing to account for multiple testing across the latent axis while maintaining sensitivity to localized expression changes.
- Statistical Rationale: The Gaussian kernel induces spatial correlation in the smoothed trajectories, implicitly modeling local dependencies in gene expression along the latent axis. By thresholding the maximum absolute deviation of the smoothed trajectory, we identify genes with sustained expression changes exceeding random fluctuations. This approach approximates a family-wise error rate (FWER) control across the latent axis, as the Gaussian smoothing reduces independent multiple comparisons.
- Empirical Validation: The threshold (1.9) was calibrated to match the 97.5% CI derived from the Gaussian null model, ensuring theoretical FPR control. Significant genes were visually highlighted (Figure 3, right), while non-DE genes were aggregated with low opacity to depict population-level variability within the CI.

## 5 Acknowledgments

We extend our sincere gratitude to the reviewers of the RECOMB 2025 conference version, as well as the additional reviewers from Cell Systems, for their insightful and constructive feedback, which substantially improved the manuscript’s quality. We also gratefully acknowledge the members of the Theis lab for their diligent proofreading and valuable suggestions, which significantly enhanced the clarity and coherence of the text. Open access funding provided by ‘Helmholtz Zentrum München— Deutsches Forschungszentrum für Gesundheit und Umwelt (GmbH)’.

## 5.1 Author contributions

K.I. conceptualized and conceived the ideas of the study, formulated the underlying hypotheses, developed the methodologies, implemented the software and the analyses, applied data preprocessing, and conducted all experimental procedures, and authored the manuscript. These were carried out under the supervision of F.J.T., who also provided guidance throughout the research process and secured funding for the project. A.R. was instrumental in the collection of metadata for human developmental datasets. A.K. contributed to the manuscript by assisting with the writing process. All authors have thoroughly read and given their approval for the final version of the manuscript.

## 5.3 Declaration of generative AI and AI-assisted technologies in the writing process

During the preparation of this work the author(s) used *ChatGPT* in order to improve language and readability. After using this tool/service, the author(s) reviewed and edited the content as needed and take(s) full responsibility for the content of the publication.

## A Supplementary Figures

**Figure 7.**
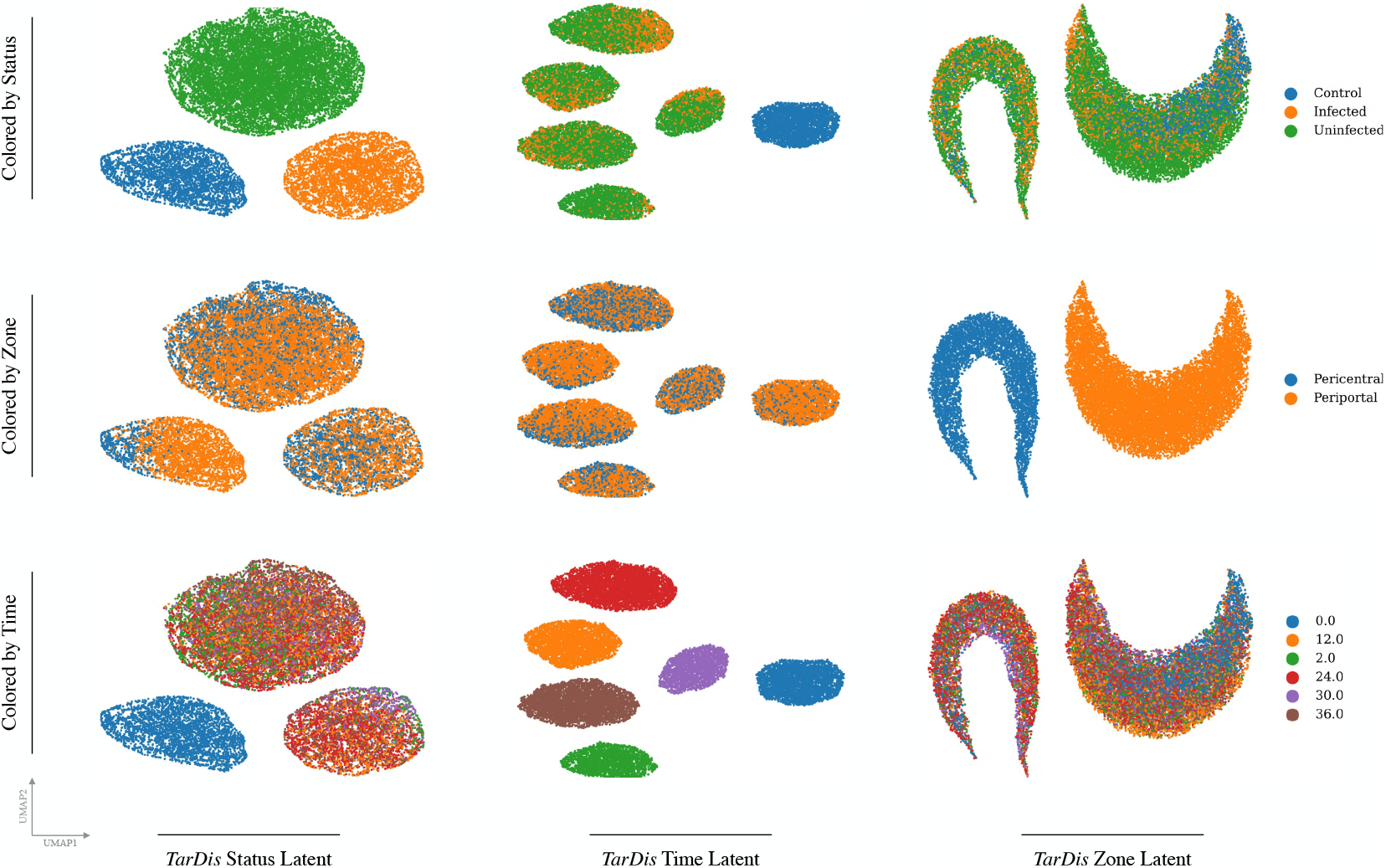
UMAP visualization of *TarDis*_multiple_ latent space representations from the *Afriat* dataset. The *TarDis* model training produces four distinct latent spaces: unreserved, status, zone, and time. The UMAP plots for the status, zone, and time latent subspaces illustrate a well-structured separation of the covariates, indicating effective encoding of the underlying data distributions and disentangled relationships within these subspaces.

**Figure 8.**
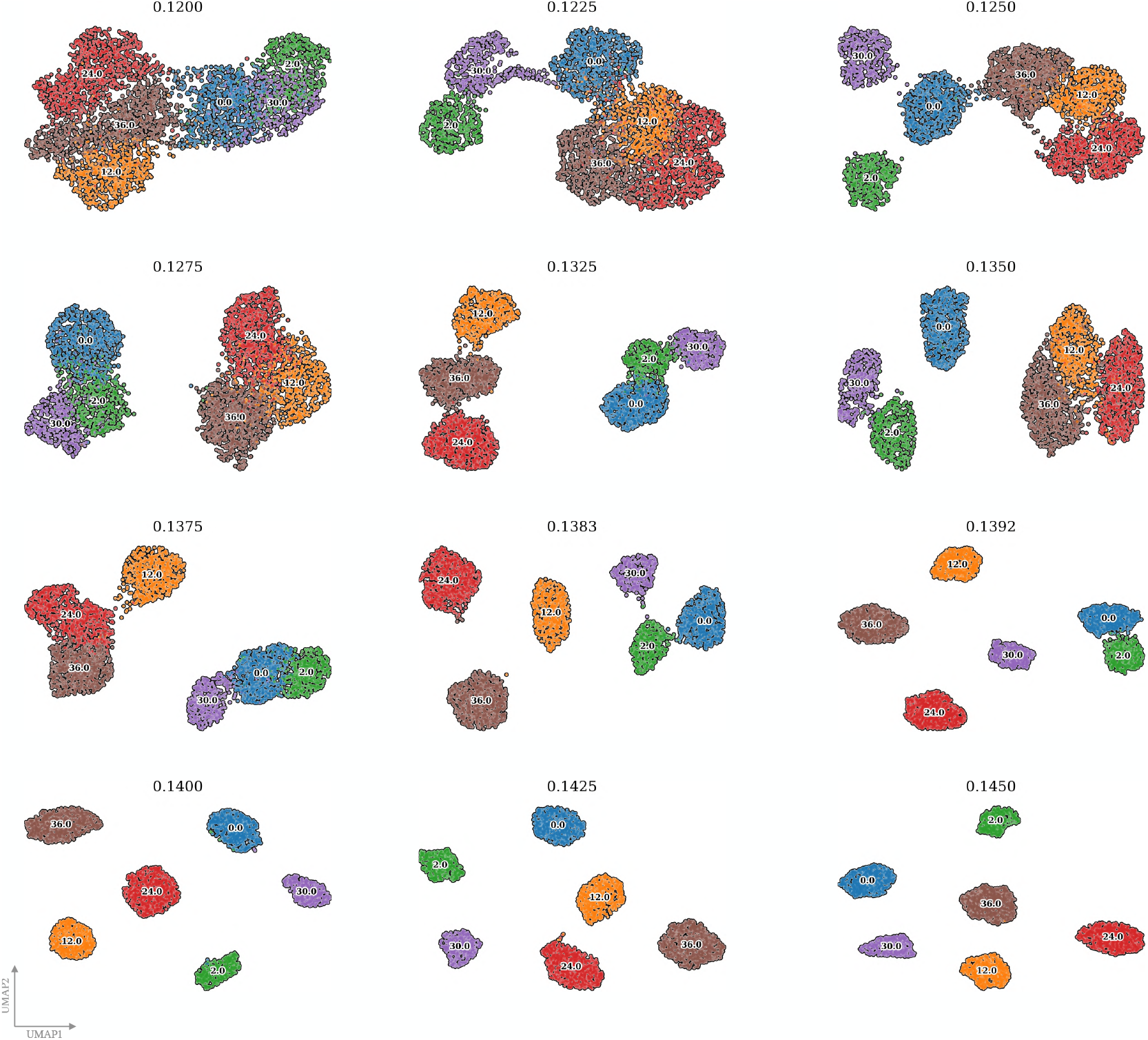
UMAP visualizations of disentangled time representation of *Afriat* dataset in the *TarDis* model with varying weights of the auxiliary loss *λ*_C_. Each panel illustrates the latent space representation of targeted time covariate, highlighting how different *λ*_C_ values influence the clustering and separation of data points corresponding to different time points. As *λ*_C_ increases, given above of the UMAP visualizations, the disentanglement quality improves, evidenced by more distinct clusters, indicating the model’s enhanced ability to preserve temporal information while disentangling other covariates. These visualizations provide qualitative support for the quantitative findings on the impact of auxiliary loss weight on disentanglement performance.

## B Supplementary Tables

**Table 6:** Complete list of differentially expressed (DE) genes identified in *Braun* and *Sciplex* datasets.

| Braun Dataset | Sciplex Dataset |
| --- | --- |
| ALKBH3-AS1 | AXL |
| ARR3 | BLOC1S2 |
| C1ORF167 | CCDC90B |
| C3ORF84 | CDKN1A |
| CD300C | DHRS7 |
| CD7 | DSC3 |
| CEBPB | FAM104A |
| CFAP141 | FAM129A |
| CNN1 | FDXR |
| EGR2 | GALE |
| EGR3 | GDF15 |
| EGR4 | GFRA1 |
| ENSG00000231977.2 | GRIP2 |
| ENSG00000258982.1 | IGFBP1 |
| ENSG00000259900.5 | MDM2 |
| ENSG00000268729.1 | MSC |
| ENSG00000272647.3 | NINJ1 |
| ENSG00000287765.1 | NKX2-8 |
| FMO10P | NPAS3 |
| FOSB | NUPR1 |
| IFT122P2 | PHLDA3 |
| KLF2 | RWDD2B |
| KLF4 | SLC52A1 |
| NR4A3 | SLFN5 |
| RTL1 | TMEM92-AS1 |
| SCX | TP53I3 |
| SPRY4-AS1 | TRANK1 |
| TMPRSS11F |  |

## Footnotes

1 *“A place for everything, and everything in its place.”* — Benjamin Franklin

2 Refer to Section 4.4.1 for discussions regarding relevant works.

3 Source code for *TarDis* is available on GitHub under `theislab/tardis`.

4 Refer to Supplementary Figure 7 for UMAP visualizations of reserved latent space representations.

5 All models were trained under configurations that aimed to closely mirror the training of *TarDis* models given in Section 4.4.3, ensuring consistency in architectural choices and the selection of analogous hyperparameters where applicable.

6 Refer to Section 4.5.2 for a comprehensive discussion on CI calculation and DE gene selection procedure.

7 Refer to Supplementary Table 6 for complete list of DE genes identified in *Braun* and *Sciplex* datasets.

8 Refer to Section 4.4.4 for a discussion of the advantages of identifying covariate-associated genes using the latent representations derived from the model, as compared to conventional differential expression analysis.

9 Refer to Section 4.4.2 for a discussion regarding the limitations.

10 Refer to Section 4.4.5 for a detailed description of the datasets used and their characteristics.

11 Refer to Figure 1b for a simplified overview of the method.

12 Refer to Section 4.5.1 for a comprehensive discussion on the different loss function options.

13 Refer to Section 4.4.3 for a detailed explanation of the chosen hyperparameters for the experiments.

14 Refer to Section 4.4.5 for a description of the datasets used and their characteristics.

15 Refer to Section 4.5.2 for a description of the evaluation metrics employed in the paper.

16 Refer to Section 4.4.4 for a discussion of the assumptions behind the theoretical framework.

